# GAGA zinc finger transcription factor searches chromatin by 1D-3D facilitated diffusion

**DOI:** 10.1101/2023.07.14.549009

**Authors:** Xinyu A. Feng, Maryam Yamadi, Yiben Fu, Kaitlin M. Ness, Celina Liu, Ishtiyaq Ahmed, Gregory D. Bowman, Margaret E. Johnson, Taekjip Ha, Carl Wu

## Abstract

To elucidate how eukaryotic sequence-specific transcription factors (TFs) search for gene targets on chromatin, we used multi-color smFRET and single-particle imaging to track the diffusion of purified GAGA-Associated Factor (GAF) on DNA and nucleosomes. Monomeric GAF DNA-binding domain (DBD) bearing one zinc finger finds its cognate site by 1D or 3D diffusion on bare DNA and rapidly slides back-and-forth between naturally clustered motifs for seconds before escape. Multimeric, full-length GAF also finds clustered motifs on DNA by 1D-3D diffusion, but remains locked on target for longer periods. Nucleosome architecture effectively blocks GAF-DBD 1D-sliding into the histone core but favors retention of GAF-DBD when targeting solvent-exposed sites by 3D-diffusion. Despite the occlusive power of nucleosomes, 1D-3D facilitated diffusion enables GAF to effectively search for clustered cognate motifs in chromatin, providing a mechanism for navigation to nucleosome and nucleosome-free sites by a member of the largest TF family.

## Introduction

In eukaryotic organisms, DNA is packaged into chromatin to regulate accessibility and recruitment of DNA-binding proteins involved in DNA replication^1^, transcription^2–4^, and genome maintenance^5^. Sequence-specific transcription factors (TFs) access specific DNA motifs (cognate motifs) on the genome to regulate transcription of target genes. Bacterial transcription factors such as LacI searches efficiently for its target on the relatively bare bacterial chromosome using a combination of 1-dimensional (1D) and 3-dimensional (3D) diffusion, referred to as facilitated diffusion^6–10^. How transcription factors search for their targets on the eukaryotic chromatin is less understood due to the packing of DNA into nucleosome particles, higher order chromatin structure and limited experimental investigations on how nucleosomes influence the search process^10^.

Chromatin can restrict accessibility of TFs at specific genomic regions by positioning nucleosomes over cognate DNA motifs^11^, thus structurally occluding half of the nucleosomal DNA surface and altering DNA conformation. Prior *in vitro* studies showed certain TFs prefer to bind at different DNA locations on a nucleosome^12–15^, and some TFs can invade nucleosomes by unwrapping DNA at the nucleosome edge or disrupting higher order chromatin organization^16–18^. Nonetheless, the search mechanisms by which any eukaryotic TF locates its targets, binds selectively to certain nucleosomal sites over others, and exhibits stable association is poorly understood. Here, we address these questions for *Drosophila melanogaster* GAF – a single-zinc-finger TF studied extensively by genetic, biochemical, genomic and live-cell, single-molecule imaging approaches^19–26^. Multi-functional GAF promotes chromatin accessibility at cognate promoter, enhancer, and insulator sites, but also in promoter-promoter and promoter-enhancer looping, assembly of the transcription preinitiation complex with the ensuing paused RNA polymerase II^27–35^. Cys_2_-His_2_ zinc finger (ZF) TFs, including GAF, constitute the largest family of eukaryotic TFs with ∼750 out of 1600 human TFs containing one or multiple ZF DNA binding domains (DBDs)^36^. The NMR structure of GAF-DBD shows the single ZF binding in the major groove to the first three bases GAG of the GAGAG consensus binding site (‘cognate motif’), and two short basic regions (BRs) N-terminal to the ZF that contact the fourth (A) and fifth (G) base in the minor and major grooves respectively^24^. Despite its single BR-ZF domain, full-length GAF (GAF-FL) forms a range of multimeric (on average hexameric) complexes through the N-terminal POZ domain^37^ and preferentially binds to clusters of cognate motifs^26,32,38^ (Fig. 1a).

**Figure 1.**
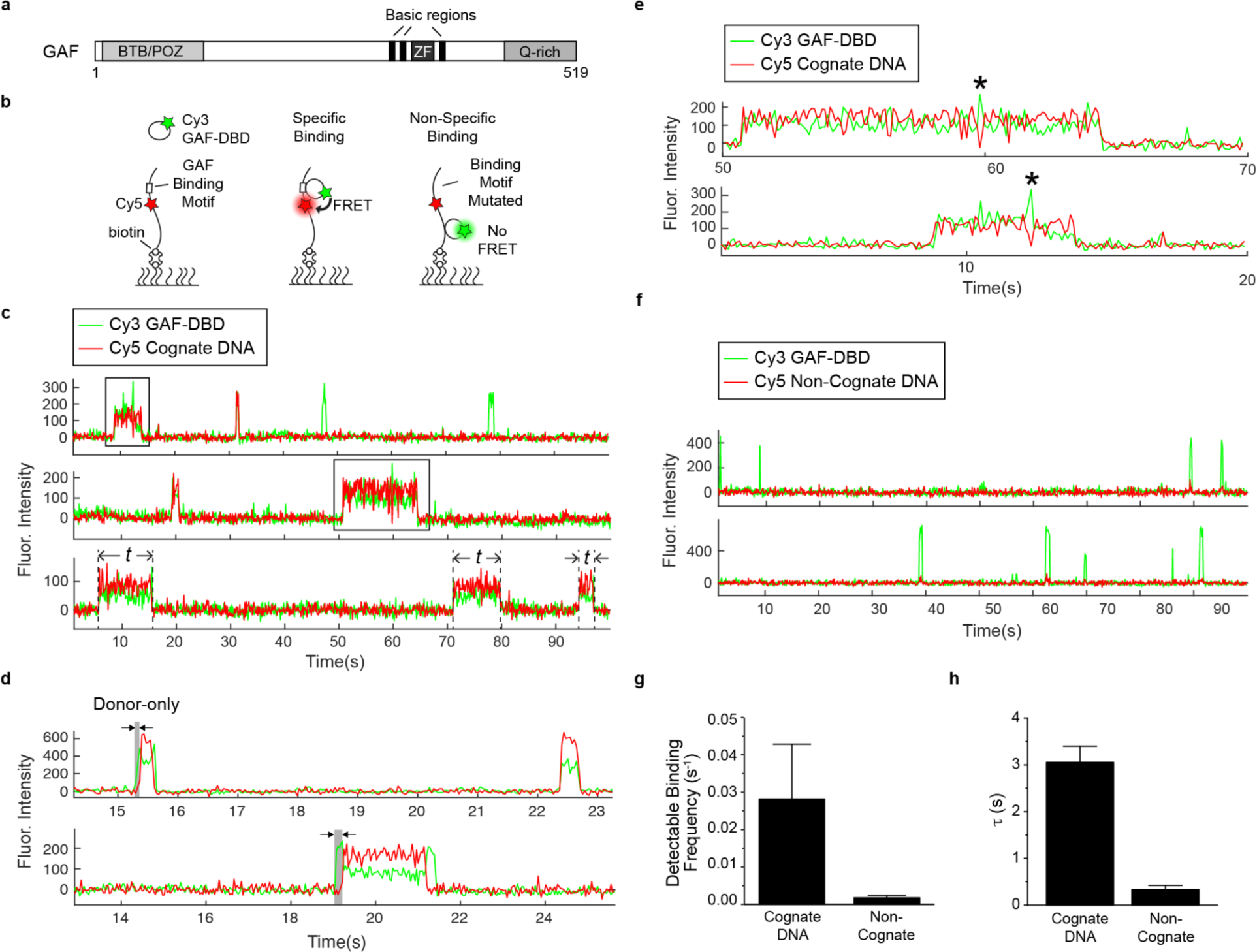
DNA sequence specificity of GAF-DBD is kinetically defined. **a**, Domain map of GAF. **b**, Schematics of two-color smFRET experiment to measure cognate-specific binding of GAF-DBD. **c**, Single-molecule trajectories showing Cy3-GAF-DBD binding to Cy5-DNA containing a GAF cognate site. Boxed binding events are zoomed in in **e**. Examples of dwell time measurements (*t*) are shown on the third trajectory. **d**, Representative single-molecule trajectories of Cy5-cognate DNA bound by Cy3-GAF-DBD showing an initial donor-only period before acceptor signal increases. The donor-only period is highlighted in grey. **e**, Zoomed-in view of binding events boxed in **c**. “*” indicates transient Cy3-only fluorescence spike during binding. **f**, Single-molecule trajectories showing Cy3-GAF-DBD nonspecifically binding to Cy5-DNA where the cognate site was substituted with a non-cognate sequence. **g**, Binding frequency of GAF-DBD to cognate or non-cognate DNA. **h**, Dwell time of GAF-DBD on cognate or non-cognate DNA. Error bars show standard deviation of three technical replicates.

Here, we used multi-color single-molecule fluorescence resonance energy transfer (smFRET)^39,40^ to study the target search process of GAF using purified components. Using FRET as a readout for cognate-motif-specific binding, we found the DNA sequence specificity of GAF-DBD is kinetically defined by both the rates of DNA association and dissociation. During target search, GAF-DBD undergoes 1D diffusion on free DNA with transient residence at cognate motifs. While remaining on free DNA, GAF-DBD escapes from the motif to locate a neighboring cognate site. Sites located at the nucleosome edge can be found by 1D invasion from linker DNA. By contrast, inner nucleosomal motifs require direct 3D association from solution but once bound, dissociation is slower. GAF-FL multimers also slide on long DNA stretched between optical tweezers and form stable complexes with naturally clustered cognate sites. Together, our results provide the first mechanistic insights on how a eukaryotic TF combines 1D and 3D diffusion to search for and locate its target sites on free DNA and nucleosomes.

## Results

### Visualizing Sequence-Specific DNA-Binding by smFRET

To investigate how GAF searches for its cognate motif on DNA, we placed a FRET donor (Cy3) at the N-terminus of GAF-DBD (by one-pot N-terminal cysteine labeling^41^; Extended Data Fig. 2) and a FRET acceptor (Cy5) on the DNA, 5 bp from a ‘GAGAGAG’ sequence consisting of two overlapping cognate motifs. The 90 bp DNA is biotinylated on one end and immobilized for smFRET imaging (Fig. 1b, left). When GAF-DBD is specifically bound to the motif, the donor-acceptor proximity results in acceptor emission via FRET (Fig. 1b, middle). When the ‘GAGAGAG’ sequence is mutated, GAF-DBD is nonspecifically bound (Fig. 1b, right). The greater average distance between the fluorophores does not allow detectable FRET, giving primarily donor emission. Under this experimental design, Cy5 fluorescence from Cy3-to-Cy5 FRET signifies motif-specific binding, while Cy3-fluorescence without FRET to Cy5 indicates nonspecific binding.

### GAF-DBD Efficiently Locates Cognate Motif on Linear DNA by 1D sliding

When surface-immobilized cognate DNA is incubated with GAF-DBD, we observe single GAF-DBD binding events as an abrupt appearance of Cy3 fluorescence above background (Fig. 1c). Motif-specific binding events show Cy5 acceptor emission during binding because of FRET (Fig. 1c). For example, in the first single-molecule fluorescence time trajectory in Fig. 1c, a GAF-DBD molecule associates specifically with the cognate motif at ∼9 s, causing acceptor emission, and dissociates at 14 s; another GAF-DBD is specifically bound at ∼31 s and quickly dissociates; two binding events at ∼48 s and 78 s show Cy3 donor emission only, demonstrating nonspecific binding. Out of all DNA binding events, GAF-DBD successfully locates the cognate motif (showing FRET) 68% ± 5% of the time. This success rate is much higher than 2% (motif length divided by entire length of DNA) – the expected probability if target search solely relies on random 3D collisions until the target motif is hit. Thus, we hypothesized GAF-DBD undergoes 1D sliding on DNA after 3D binding to search for its specific target (hereafter, 1D sliding is used interchangeably with 1D diffusion and includes both helically-coupled 1D sliding as well as 1D hopping). Accordingly, a time lag between the appearance of donor and ensuing acceptor fluorescence represents the period between GAF-DBD landing anywhere on the DNA and successful recognition of the target motif. For most (∼80%) binding events, the time delay is shorter than instrument resolution (35 ms), but we did occasionally detect such delays (Fig. 1d). In addition, we observe transient Cy3 donor-only fluorescence flanked FRET events, which suggests GAF-DBD transiently slides off the target quickly before returning (Fig. 1e, asterisks). At lower ionic strength, the nonspecific dwell time increases, suggesting 1D diffusion involves some 1D hopping along the DNA^42^; the longer dwell time at nonspecific sites was also consistent with documentation that TFs can lose sequence-specificity under low-salt^43^ (Extended Data Fig. 3).

### DNA Binding Kinetics Defines Sequence Specificity of GAF-DBD

To compare sequence-specific to nonspecific DNA binding, we examined GAF-DBD kinetics on non-cognate DNA where GAGAGAG was replaced by TATACAG, observing only transient Cy3 donor emissions (Fig. 1f). Because of this complete absence of FRET signal, we could reliably use FRET appearance as a reporter for motif-specific binding. At the same GAF-DBD concentration, detectable binding attempts on non-cognate DNA (0.0018 s^-1^ or 1 binding attempt every 560 s) are 16-fold less frequent than on cognate DNA (0.028 s^-1^ or 1 binding attempt every 36 s; Fig. 1g). The GAF-DBD dwell time on non-cognate DNA (1/*k*_off_ = 0.33 s; time spent on DNA, either immobile or 1D sliding) is 9-fold less than on cognate DNA (3.1 s; Fig. 1h). Therefore, the cognate site both increases binding frequency (*k*_on_) and transiently traps GAF-DBD to reduce dissociation (*k*_off_) achieving high overall sequence-specificity. These results demonstrate sequence-specific binding of GAF-DBD is defined by both association and dissociation kinetics.

### Three-Color FRET Reveals 1D Back-and-Forth Sliding of GAF-DBD

On the *Drosophila* genome, GAF preferentially binds to closely spaced cognate motifs at numerous promoters and enhancers^38^. To mimic a native GAF target, we used a 187-bp segment from the *Drosophila hsp70* promoter sequence, retaining the two longest GAF binding sites while several other short GA-elements were mutated (Fig. 2a). Two FRET acceptors for the Cy3 donor on GAF-DBD, Cy7 and Cy5, were placed 3 bp away from Site 1 ‘GAGAGGGAGAGA’ and 5 bp away from Site 2 ‘GAGAGAG’ respectively with 57 bp of intervening DNA. FRET detection signifies GAF-DBD binding to the corresponding sites, while Cy3 emission without FRET reports nonspecific binding (Fig. 2a).

**Figure 2.**
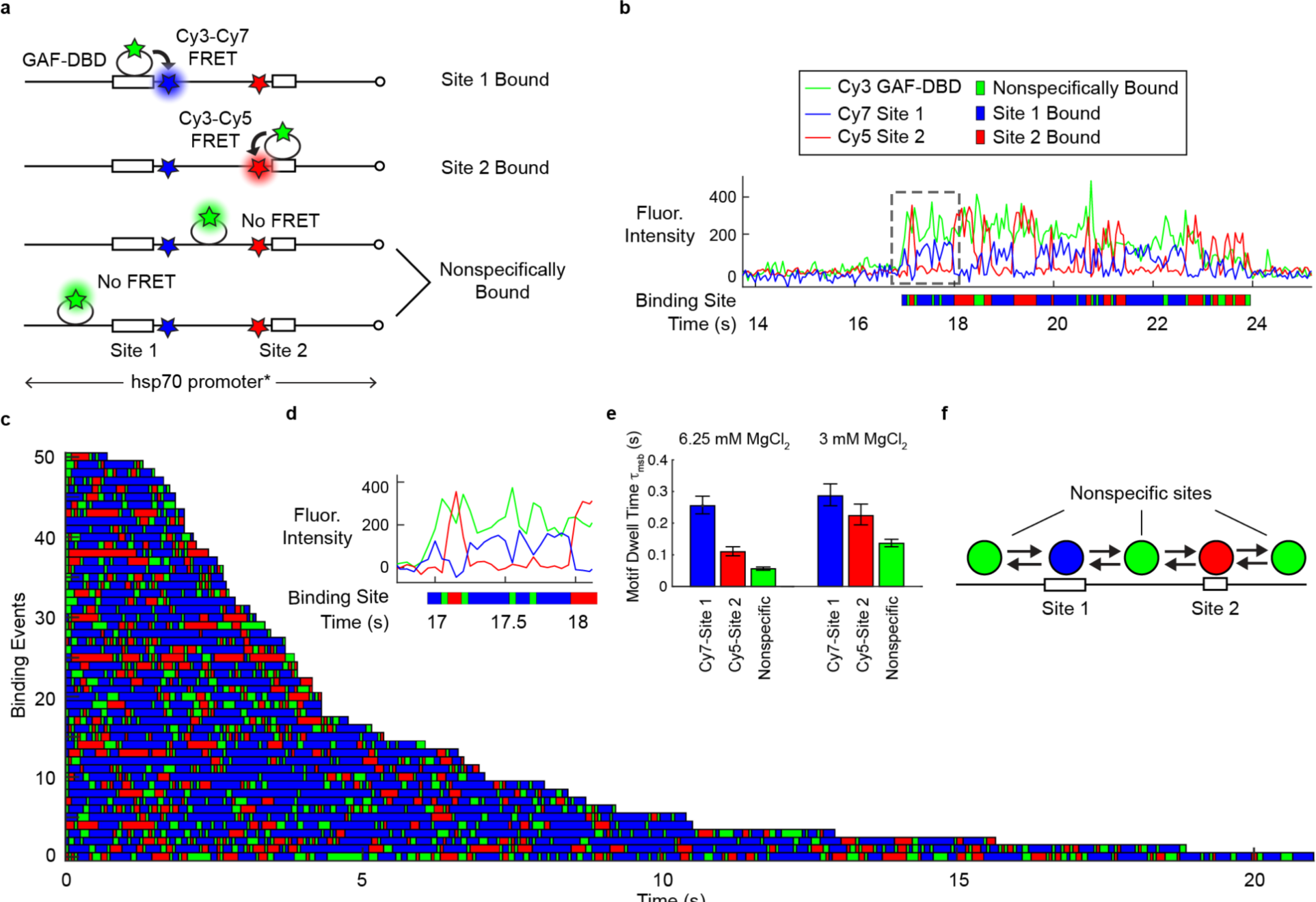
GAF-DBD explores free DNA by 1-dimensional diffusion. **a**, 3-color smFRET experiment distinguishes whether GAF-DBD is bound to Cy7-labeled Site 1, Cy5-labeled Site 2, or a nonspecific site on *Drosophila* hsp70 promoter DNA. **b**, A single-molecule trajectory shows GAF-DBD sliding back-and-forth on the DNA between two cognate motifs. Binding site assignment is shown below as a colored ribbon. **c**, 55 binding events shown as a rastergram **d**, Zoomed-in view of the boxed region in **b**. **e**, GAF-DBD dwell times on Site 1, Site 2, or nonspecific site on the DNA at regular (6.25 mM) or low (3 mM) MgCl_2_. **f**, Schematic of GAF-DBD sliding on DNA. Circles represent GAF-DBD. Green circles indicate GAF-DBD binding to a nonspecific site; blue circle, GAF-DBD binding to Site 1; red circle, GAF-DBD binding to Site 2.

In the single-molecule trajectory of Fig. 2b (expanded in Fig. 2d), a GAF-DBD molecule lands on the DNA at ∼17 s, emitting Cy3 fluorescence. Almost immediately, we observe Cy3-Cy7 FRET, suggesting GAF-DBD locates Site 1. Cy7 intensity drops and Cy5 intensity increases 0.25 s later, indicating GAF-DBD slides away from Site 1 to reach Site 2, emitting only Cy3 fluorescence during transit. 0.25 s later, GAF-DBD slides back to re-engage with Site 1, exciting the Cy7 fluorophore. As noted, we also observe intervening periods of Cy3 fluorescence with neither Cy5-nor Cy7-FRET, indicating nonspecific binding during transit. We interpret these FRET behaviors as GAF-DBD undergoing 1D diffusion over the entire DNA fragment.

To automate assignment of binding sites for these trajectories, we developed a Hidden Markov model (HMM) that uses 3-color fluorescence intensities to infer a most likely sequence of GAF-DBD binding positions (Supplementary Note). The results are plotted as colored ribbons where a blue segment represents Cy7 Site 1 binding; red, Cy5 Site 2 binding; and green, nonspecific binding (Fig. 2b); ribbon length indicates the duration of binding. For all 55 randomly selected ribbons stacked to form a rastergram, blue and red segments are interspersed with green, indicating GAF-DBD undergoes 1D diffusion for every specific binding event (Fig. 2c). On average, dwell times on Site 1 (blue, τ_1_ = 0.25 s) are longer than Site 2 (red, τ_2_ = 0.11 s) (Fig. 2e), likely because Site 1 (12 bp) has a more extended cognate site than Site 2 (7 bp). The overall dwell time on this DNA fragment is 3.34 +/- 0.08 s, similar to the 3 s dwell time on the 90 bp DNA harboring a single binding site with the same sequence as Site 2 (Fig. 1g).

At lower ionic strength (3 mM versus 6.25 mM MgCl_2_), GAF-DBD exhibits a longer dwell time at non-specific sites (Cy3-only state lifetime is 0.14 s versus 0.056 s, respectively) (Fig. 2e, Extended Data Fig. 4a, b), consistent with observations on the single-motif DNA (Extended Data Fig. 3). When Site 1 (Cy7) is mutated, Cy3-Cy7 FRET becomes transient or undetectable, as expected, while Cy3-Cy5 FRET remains stable (Extended Data Fig. 4c-e). Therefore, alternating FRET between Sites 1 and 2 is a consequence of 1D sliding back and forth on DNA rather than artifactual fluorophore binding or conformational changes of the DBD itself. Hence, we conclude GAF-DBD undergoes 1D diffusion on DNA during its search (Fig. 2f), and importantly, with a functional outcome for target location.

On the natural *hsp70* promoter, Site 1 and Site 2 are located on different strands of the DNA. To reflect the natural orientation, we replaced Site 1 with the complementary sequence ‘CTCTCCCTCTCT’. Thus, the GAF ZF must re-orient 180 degrees orthogonal to the DNA axis for motif recognition. Interestingly, GAF-DBD still slides between inverted Site 1 and Site 2 (Extended Data Fig. 4f-h) with a similar overall dwell time on DNA (3.6 +/- 0.05 s compared to 3.34 +/- 0.08 s), indicating that GAF-DBD can readily flip on DNA during 1D diffusion to accommodate opposite motif orientations.

### GAF-DBD Also Directly Binds to Sites from 3D

Given that GAF-DBD slides between neighboring binding sites for every specific binding event, we wondered whether 1D sliding always precedes a target strike, as shown by a delay between nonspecific DNA binding (Cy3 only) and site recognition (FRET). We observed such delay for 40-50% of binding events; the remainder displays FRET immediately upon GAF-DBD landing, i.e. binding by 3D diffusion (Extended Data Fig. 4i). While camera exposure time (50 ms) would not resolve 1D diffusion from a nonspecific site very close to target, at lower ionic strength (MgCl_2_ reduced from 6.25 mM to 3.00 mM) longer residence (136 ms) on nonspecific sites should reveal missed sliding events; yet we still observe FRET at 39% frequency upon binding (Extended Data Fig. 4i), suggesting that most immediate FRET-upon-binding events can be ascribed to direct binding from 3D.

### Nucleosome Impedes 1D Sliding into its Core

We next asked whether GAF-DBD uses 1D diffusion to find cognate motifs within a nucleosome. We tested three 40-N-40 mono-nucleosome constructs, each containing a Cy5-labeled ‘GAGAGA’ site on the linker DNA and a second Cy7-labeled ‘GAGAGA’ site at distinct superhelical locations SHL7, SHL5 or SHL3 inside a nucleosome positioned by the Widom 601 sequence^44^ (the major groove of each motif facing the histone core) (Fig. 3a). For the nucleosome bearing a motif at SHL7, we observe alternating Cy5 and Cy7 fluorescence, indicating 1D sliding between linker and SHL7 sites (Fig. 3b, e). 75% of 485 binding events shows sliding between the SHL7 site and linker DNA, 19% shows stable residence on the linker DNA site, and 7% reside on the SHL7 site only (Fig. 3h). In contrast, sites placed at SHL5 or SHL3 are rarely bound: ∼90% of binding events show Cy3-Cy5 FRET, suggesting that binding to the linker DNA site dominates over SHL5 and SHL3 sites (Fig. 3c, d, f, g, h).

**Figure 3.**
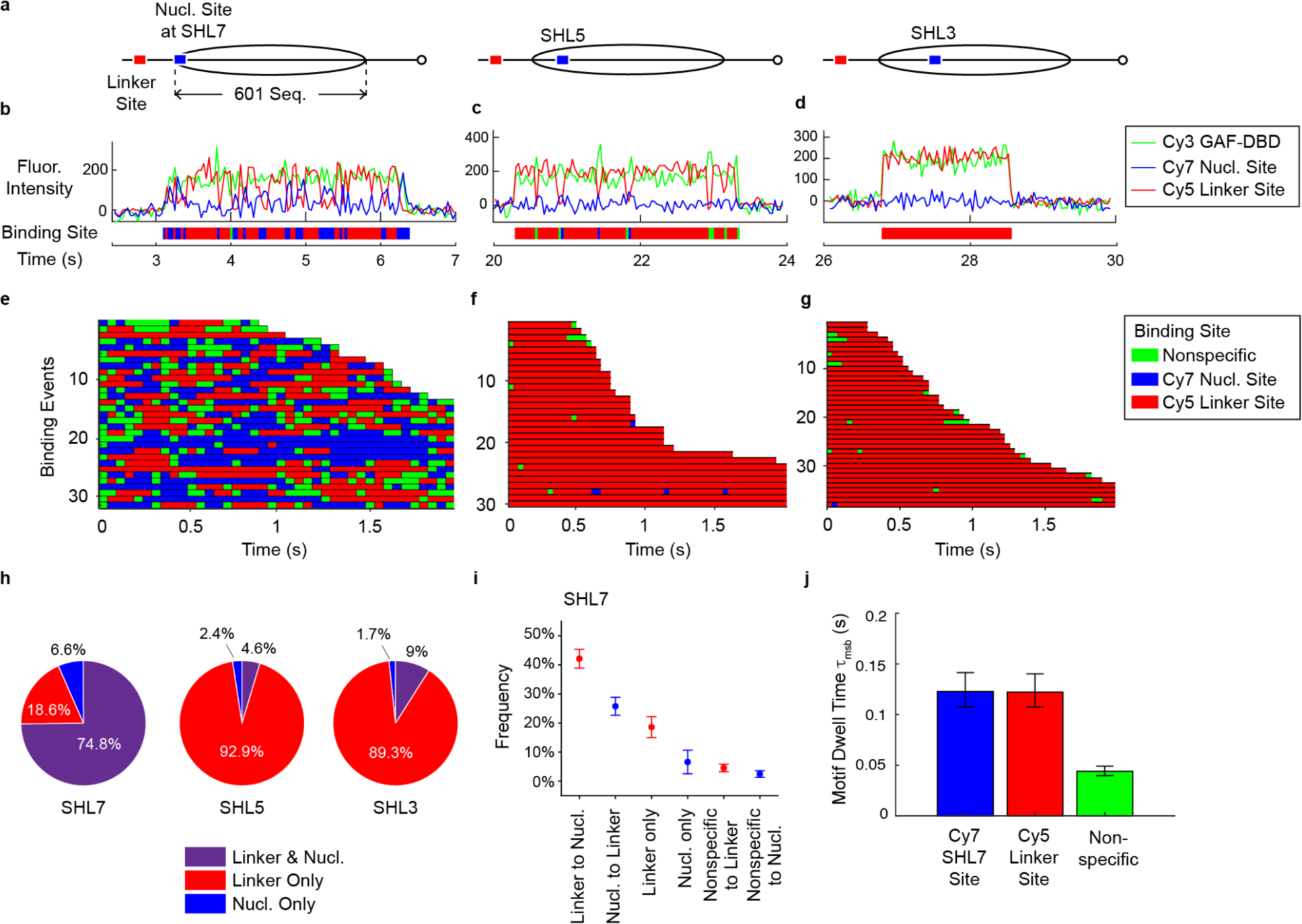
Nucleosome blocks GAF-DBD 1D sliding beyond SHL7. **a**, Schematics of nucleosome constructs where the nucleosomal Cy7-labeled cognate site (blue) is placed at SHL7, SHL5, or SHL3 locations and the other Cy5-labeled cognate site (red) is on linker DNA. The nucleosome is positioned by the Widom 601 sequence flanked by 40 bp linker DNA on both sides (40-N-40). **b-d**, Representative single-molecule trajectories of GAF-DBD binding to linker or nucleosomal sites at SHL7 (**b**), SHL5 (**c**) and SHL3. **e-g**, Binding site rastergrams for GAF-DBD on nucleosome constructs where the Cy7-labeled motif is located at SHL7 (**e**), SHL5 (**f**) or SHL3 (**g**). **h**, GAF-DBD binding categories for each construct. N=485 for SHL7, N=260 for SHL5, N=272 for SHL3. **i**, Categories of binding events on SHL7 construct. **j**, GAF-DBD motif dwell times on SHL7, linker site and nonspecific sites on the SHL7 construct.

To narrow down the nucleosome boundary that blocks 1D sliding, we measured nucleosome binding for a cognate site at SHL6.5. Binding to this nucleosome resembles SHL3- and SHL5-site nucleosomes, with much fewer sliding events (4.5%) and more binding to linker site only (83.3%) compared to the SHL7-site construct (67.2% sliding, 19.7% linker site binding only) (Extended Data Fig. 5). These results indicate nucleosome organization allows 1D sliding of GAF-DBD to no further than its outer edge, effectively blocking sliding even 5 bp deeper into the nucleosome core. Thus, transient, intrinsic DNA-unwrapping dynamics^45^ is apparently insufficient for 1D sliding into the nucleosome core.

To bypass this blockade, GAF-DBD could bind directly by 3D diffusion to solvent-exposed nucleosome locations. As the major groove of the cognate sites at SHL5 and SHL3 face the histone core, we hypothesized that shifting the cognate sites by 5 bp each to SHL4.5 and SHL2.5 to face the solvent should result in a higher fraction of direct 3D binding to the nucleosomal site (stable Cy3-Cy7 FRET throughout binding). Indeed, the SHL4.5 site exhibits a 12-fold higher fraction (22.6%) of nucleosome binding via 3D diffusion than the SHL5 site (1.9%), and the SHL2.5 site exhibits a 3-fold increase (20.9%) compared to SHL3 (7.1%) (Extended Data Fig. 5). These results demonstrate solvent-facing cognate motifs are indeed more accessible by 3D diffusion.

### Linker DNA is the Preferred Landing Site over Nucleosome Core

Among binding events leading to sliding to the SHL7 site, 62% initially lands on the 40 bp linker DNA site, compared to 38% directly binding to SHL7 (Fig. 3i,), indicating 1.6-fold greater accessibility (association rate) for a site on the nucleosome linker than on its edge. Less common is binding to linker DNA site only (19%, e.g., Fig. 2e, row 3, constant red), but still more frequent than 3D binding to SHL7 site only (7%; constant blue), a further indication that the linker site is more accessible (Fig. 3i). Occasionally, a Cy3-only state appears before FRET to Cy5 or Cy7 (‘Nonspecific to Linker’ or ‘Nonspecific to Nuc’), corresponding to the aforementioned delay between initial nonspecific binding and target engagement. We also observe that, on average, GAF-DBD has similar dwell times (τ_msb_) on the linker (0.12 s) and SHL7 (0.12 s) sites, indicating nucleosome edge location does not affect the dissociation rate (1/τ_msb_) for GAF-DBD (Fig. 3j). Hence, location of a cognate site on the nucleosome edge mildly reduces the on-rate but does not affect the off-rate for GAF-DBD binding.

### 3D Diffusion Enables Nucleosome Target Search

On native promoters such as *Drosophila hsp70*, several GAGA elements over multiple phases along the DNA helical axis^23^ could potentially increase 3D accessibility. Accordingly, we designed and constructed by linker ligation^46^, a 0-N-40 *hsp70* promoter nucleosome whose position (–209 to –63) mimics one of three thermodynamically favored nucleosomes on hsp70 promoter DNA harboring two native GAGA elements^47,48^. As shown in Fig. 4a, the ‘CTCTCCCTCTCT’ site near the nucleosome dyad is labeled with Cy7, while the other, ‘GAGAGAG’ at the nucleosome edge is Cy5-labeled. We hypothesized that GAF-DBD should locate the nucleosome dyad site predominantly through 3D diffusion, while 1D or 3D diffusion should find the site at the nucleosome edge. Fig. 4b shows idealized traces of GAF-DBD searching for the dyad site by 3D diffusion: the first binding event directly targets the Cy7-dyad site, exhibiting only Cy3-Cy7 FRET throughout the duration of binding; the second event occurs at the Cy5-edge site via 1D or 3D diffusion, showing only Cy3-Cy5 FRET. We observe three types of binding events (N=160): a) GAF-DBD binding to the edge site (48%, Fig. 4c); b) to the dyad site distal to Cy7, with FRET to both Cy5 and Cy7 acceptors (36%, Fig. 4d); c) to the dyad site proximal to Cy7, with FRET to Cy7 only (13%, Fig. 4e). There is also residual binding to un-reconstituted free DNA showing alternating FRET to Cy5 or Cy7 (3%, Fig. 2b). Remarkably, 97% of binding events reveal no anticorrelated changes in Cy5 and Cy7 FRET compared to binding to free DNA, indicating that GAF-DBD remains bound to one site without sliding or hopping to the other while remaining on the nucleosome (Fig. 4f). Taken together, we find that 3D diffusion is the major search mode for locating inner nucleosomal motifs.

**Figure 4.**
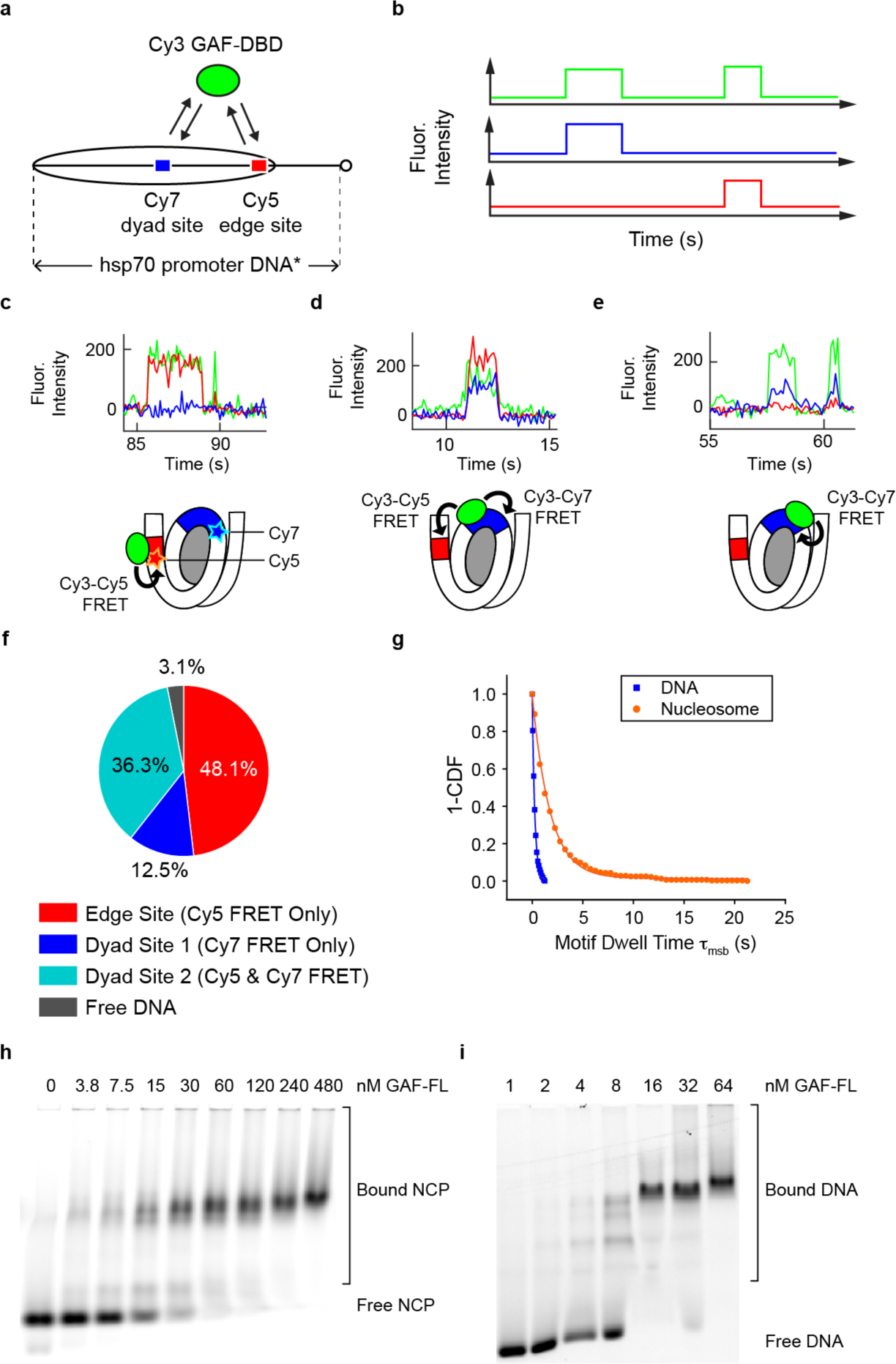
3D diffusion dominates for inner nucleosomal targets. **a**, *Drosophila hsp70* promoter nucleosome construct (0-N-40) for investigating GAF-DBD search when two binding sites are within nucleosome core. **b**, Idealized single-molecule trajectories for GAF-DBD locating two nearby cognate sites. **c-e**, Representative single-molecule trajectories on this nucleosome construct. 0.4 nM Cy3-GAF-DBD was used for these experiments. **f**, Binding event categories on the nucleosome. **g**, 1-CDF of motif dwell times for nucleosome (orange) compared with free DNA (blue). Free DNA data are from Cy7-Site 1 dwell time (Fig. 2e, 6.25 mM MgCl_2_). **h**, EMSA showing GAF-FL binding to Cy5-NCP formed on *hsp70* promoter DNA fragment. **i**, EMSA showing GAF-FL binding to Cy5-DNA. The same DNA was used to reconstitute Cy5-NCP in **h**.

Interestingly, individual nucleosomes exhibit sustained preference for one 3D-binding location over others (Extended Data Fig. 6a). After initial binding to and dissociation from the nucleosome edge (Cy3-Cy5 FRET) (type A), frequent re-binding (88%) occurs at the same location (type A-A; Extended Data Fig. 6b). The same re-binding preference is observed for type B-B (84%) and type C-C events (88%), suggesting that the reconstituted *hsp70* nucleosome population is represented by at least three distinctly phased, stable configurations, each presenting a different preferred site for 3D binding. We occasionally observe transient Cy3 fluorescence without FRET for the duration of site-specific binding on the nucleosome (Extended Data Fig. 6: black asterisks), suggesting that GAF-DBD may undergo ultra-short range (<10 bp) 1D diffusion within the exposed cognate site. The motif-specific dwell times of GAF-DBD on the nucleosome are strikingly 10-fold greater than on naked DNA motifs (Fig. 4g). This indicates that nucleosome architecture can better trap GAF-DBD at exposed cognate sites on the core histone surface than free DNA, reducing the rate of dissociation and providing some compensation for the nucleosomal impediment to 1D and 3D association.

GAF-FL consists of an N-terminal POZ multimerization domain, a central DBD and a C-terminal Q-rich region (Extended Data Fig. 2b). Biochemical studies^37^ show GAF-FL forms multimeric complexes (on average, hexamers) via the POZ domain and binds preferentially to a natural cluster of cognate sites (as found on the *hsp70* promoter, Extended Data Fig. 2c). Our results with GAF-DBD suggest that full-length multimers should also directly bind to (linker-free) nucleosome core particles (NCPs) if cognate motifs are solvent-exposed. Accordingly, we performed electrophoretic mobility shift assay (EMSA) on GAF-FL and (3-N-6) *hsp70* NCP binding. The results show that GAF-FL binds less well to the NCP than to bare DNA but only by ∼2-fold (compare bound fractions in the titration series, Extended Data Figure 6). Since the reconstituted NCP contains almost no linker DNA for 1D sliding, we infer that the cognate sites must be found by 3D diffusion for GAF-FL.

### Full-Length GAF Displays 1D Sliding Before Locking on Hsp70 Target

To investigate whether GAF-FL also undergoes 1D diffusion on DNA, we tracked the position of SNAP-tagged GAF-FL labeled with AlexaFluor488 (Extended Data Fig. 7) on kilobase pairs (kbp) long DNA stretched between two optical traps (C-Trap, LUMICKS; Fig. 5b, Extended Data Fig. 8c). This DNA is a concatenated 3.3 kbp plasmid carrying a 359-bp insert containing the *hsp70* promoter harboring a dense cluster of full and partial GAGAG sequences (vector+*hsp70)* (Fig. 5a, Extended Data Fig. 8a,b). Accordingly, adjacent promoters can be about 0, 1 or 2 plasmid lengths apart, giving rise to a semi-random distribution of *hsp70* promoters over tens of kbp DNA.

**Figure 5.**
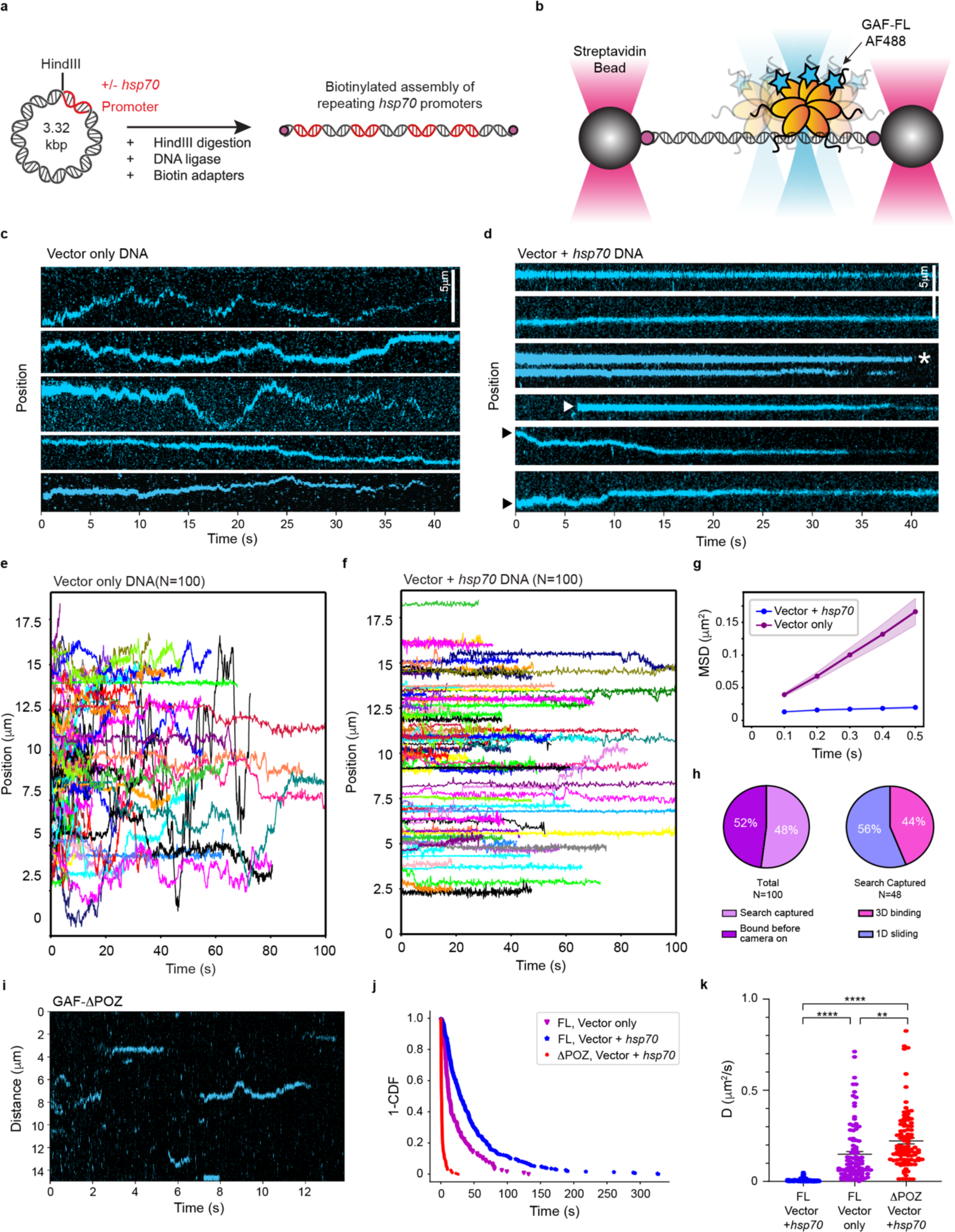
Full-length GAF multimer undergoes 1D diffusion during target search. **a,** Schematic of construct design. Plasmids with or without the *hsp70* promoter sequence were digested with HindIII and concatenated using T4 DNA ligase. Exposed ends were then biotinylated with adaptors, resulting in dual-end biotinylated DNA carrying repeating *hsp70* promoters. **b**, Set-up of dual optical tweezers for confocal microscopy of stretched DNA tethered between two streptavidin-coated polystyrene beads. **c-d**, Representative kymographs show AF488-GAF-FL fluorescence signal (cyan) on DNA over time in the absence (c, vector only) or presence of *hsp70* promoter (d, vector+*hsp70*). White arrowhead shows GAF-FL binding directly to target from 3D; black arrowheads show GAF-FL undergoing 1D search before finding target; white asterisk shows direct 3D dissociation. **e,** Compiled plots of position versus time for GAF-FL trajectories on vector only DNA. 100 traces were collected for each condition and arranged to start from zero time. **f**, Compiled plot for GAF-FL positions over time on vector+*hsp70* DNA. **g**, Average mean squared displacement (MSD) over time lag of all collected traces for vector only (purple) and vector+*hsp70* DNA (blue). Shaded area reports standard error of the mean (SEM). N=100. **h**, Pie charts categorizing GAF-FL traces by DNA binding at onset of movie (left), and targeting directly (3D binding) or indirectly (1D sliding) (right). **i**, A representative kymograph of GAF-ΔPOZ on vector+*hsp70* DNA. **j,** 1-CDF comparing dwell times of GAF-ΔPOZ on vector+*hsp70* DNA (red, τ = 1.67 s), GAF-FL on vector+*hsp70* (blue, τ = 43.4 s) and GAF-FL on vector-only (purple, τ = 21.0 s). **k**, Single-trace diffusion coefficients for GAF-FL on vector+*hsp70* and vector-only DNA and GAF-ΔPOZ on vector+*hsp70* DNA. Error bars are SEM. Statistical differences were determined by unpaired t-test (n=100, **** = p < 0.0001).

On vector only DNA, multimeric GAF-FL displays rapid, random 1D diffusion with a mean D_coef_ = 0.149 μm^2^/s or 1606 bp^2^/s, often traversing a few μm (∼10 kbp) per minute (Fig. 5c, e). A minuscule fraction (4%) shows non-diffusive binding (D_coef_ < 0.01 μm^2^/s), likely because of full or partial motifs in isolation on the vector. In contrast, on vector+*hsp70* DNA, GAF-FL shows mostly (78%) non-diffusive binding and remains immobile for at least 40 s before photobleaching or dissociation (Fig. 5d, f). The average mean squared displacement (MSD) over time lag quantifies the different mobility of GAF-FL on vector+*hsp70* versus vector only DNA (Fig. 5g). Although a substantial fraction of GAF-FL has immobilized on target between protein injection and imaging start (∼30 s), 44% of *de novo* binding events appear abruptly and remain immobile, indicative of 3D targeting (Fig. 5h; Fig. 5d, white arrowhead; Extended Data Fig. 8e), while 56% exhibit detectable 1D diffusion before stasis, consistent with a 1D search after stochastic 3D nonspecific binding (Fig. 5d, black arrowhead; Extended Data Fig. 8d). Dissociation of GAF-FL from target also occurs by 1D where GAF-FL slides off target to re-initiate a 1D search (34%, Extended Data Fig. 8f), or by 3D where it dissociates into solution without return (66%; Fig. 5d, white asterisk).

Calculation of the 1D D_coef_ for all GAF-FL traces revealed a wide distribution of D_coef_ values on vector DNA (Fig. 5k; vector only), likely because of heterogeneity in multimeric states (mass differences) and also in the number of DBD-DNA interactions within each GAF-FL multimer. The multimeric state of GAF-FL is also consistent with our observation of the gradual loss of AF488 fluorescence from photobleaching during DNA binding (Extended Data Fig. 9d, d). To examine the properties of monomeric GAF, we tracked GAF deleted for the POZ domain (GAF-ΔPOZ, which retains the DBD and Q-rich region; Extended Data Fig. 7d) and found that DNA-bound GAF-ΔPOZ undergoes single-step photobleaching, indicative of its monomeric state (Extended Data Fig. 9b, e). We were able to distinguish GAF-ΔPOZ photobleaching events from dissociation events (Extended Data Fig. 9c).

Moreover, GAF-ΔPOZ monomers showed strikingly transient binding (1.67 s) on vector+*hsp70* DNA (Fig. 5i, j, Extended Data Fig. 9a) compared to GAF-FL (43 s). This transience is similar to the 3-4 s dwell time of monomeric GAF-DBD on short DNA fragments mentioned above. In addition, GAF-ΔPOZ D_coef_ on vector+*hsp70* DNA is 31-fold higher than GAF-FL (0.222 vs 0.007 μm^2^/s) and even slightly higher than GAF-FL on nonspecific DNA (0.149 μm^2^/s) (Fig. 5k). This indicates that multimerization via the POZ domain is necessary for site-specific binding *in vitro* at timescales approaching in vivo values (∼2 min)^26^. Overall, these results show that specific and prolonged GAF-FL binding on a physiological target sequence requires both protein multimerization and clustered cognate sites.

## Discussion

3-color FRET is an increasingly popular approach for studying complex DNA processes, including LacI target search^9^, chromatin remodeling^49,50^,and DNA repair^51^. Inspired by the study on LacI’s search mode for the Lac operator by 1D-3D facilitated diffusion on bacterial DNA^9^, we used multi-color FRET to explore whether a eukaryotic TF relies on 1D diffusion for target search on free DNA and a nucleosomal template in which histone-DNA interactions dominate overall DNA accessibility. Our work reveals that the ZF DBD of GAF searches for its cognate DNA and nucleosome targets by a combination of 1D and 3D diffusion. 1D diffusion occurs back and forth over several hundred bp as the major search mode to locate cognate motifs on nucleosome-free DNA, similar to LacI^9^. In addition, 1D sliding can also penetrate into the accessible edge (SHL7) of nucleosomes, likely because of the transient, intrinsic unwrapping of the entry-exit sites of nucleosomal DNA^52,53^ (Extended Data Fig. 1a). Using optical tweezers and single particle imaging, we further showed that multimeric GAF-FL harboring multiple ZF DBDs also uses 1D diffusion to search for natural promoter targets on naked DNA (Extended Data Fig. 1b). The POZ multimerization domain is required for maintaining DNA interactions during this 1D search and, once the target of clustered cognate sites is located, GAF-FL remains on-site for near physiological time periods.

However, 1D diffusion cannot reach motifs at SHL 6.5 or locations deeper into the nucleosome core, rendering this search mode ineffective at locating inner nucleosomal sites. The second search mode, 3D diffusion, allows GAF-DBD or GAF-FL not only to target sites on naked DNA, but also to find solvent-facing, but not core histone-facing, nucleosomal motifs by direct association from solution (Extended Data Fig. 1a). The phasing of the two cognate sites on the reconstituted *hsp70* promoter nucleosome^13,15^ are such that the major groove of at least one motif faces solvent. In addition, detection of ultra-short range (single bp-scale) intrinsic DNA mobility of the native *hsp70* sequence relative to the histone octamer may allow different rotational phases to be sampled for better motif recognition^54,55^. Thus, on eukaryotic chromatin, GAF would combine 1D sliding for accessible linker and nucleosome-free DNAs, and 3D diffusion for targeting exposed motifs on core nucleosomes, achieving specificity for nucleosomal and nucleosome-free sites through differential binding kinetics.

GAF-FL multimers form via the POZ domain and each multimer therefore contains multiple DBDs. Our long-range measurements (C-Trap) show that GAF-FL also undergoes 1D diffusion on DNA. In good agreement with findings for GAF-DBD, GAF-FL apparently locates a substantial portion of binding sites directly by 3D diffusion (although short-range 1D sliding beyond instrument resolution from a nearby nonspecific site would be mis-counted as 3D binding). Monomeric GAF-ΔPOZ also exhibits robust 1D sliding, indicating that multimerization is not required for 1D diffusion. However, GAF-ΔPOZ shows poor site specificity as its diffusion coefficient on target-containing DNA is even higher than GAF-FL on nonspecific DNA, indicating that without the POZ domain, GAF cannot remain stably associated with its cognate site.

The stably bound GAF-FL population resides on chromatin for ∼2 min in living *Drosophila* hemocytes, likely at cognate promoter/enhancer NDRs^26^. Consistent with this, purified GAF-FL remains on free *hsp70* target DNA stretched between optical tweezers for a minimum of 40 s *in vitro*. Multimerization is necessary for stable association *in vitro*, as removal of the POZ domain dramatically reduces GAF dwell time on DNA to ∼2 s. This difference can be explained if GAF-FL dissociation requires multiple DBDs to dissociate from all engaged motifs – a low probability event compared to a single DBD dissociating from one site. Interestingly, the *in vivo* residence time of the transiently bound GAF-FL population (3.7 s)^26^ is similar to the *in vitro* specific dwell time of GAF-DBD and GAF-ΔPOZ, suggesting that transient binding *in vivo* may be due not only to GAF-FL multimer binding nonspecifically, but also, as this study suggests, to specific binding between one DBD (in monomeric or multimeric GAF) and an isolated cognate motif.

Prior evidence has established that GAF has a key function to establish and maintain (or to ‘pioneer’) nucleosome-depleted regions in chromatin: 1) GAF is necessary for the creation of accessible chromatin at inducible, developmental and house-keeping *Drosophila* gene promoters *in vivo* and *in vitro^23,29^*; 2) it directly recruits ATP-dependent chromatin remodelers to generate DNaseI-hypersensitive sites^28,56^; and 3) it exhibits an uncommonly long residence time and high site occupancy from live-cell single-molecule imaging studies^26^. We find that the target search mode of GAF *in vitro* is highly influenced by the chromatin environment. The impediment of nucleosome organization to 1D sliding implies that a 1D search could start anywhere on nucleosome-depleted DNA with site targeting after rapid back and forth 1D sliding, while nucleosomal targets must be accessed by 3D diffusion, and only if sites are rotationally phased to solvent exposure. Accordingly, GAF and other TFs have a lower *k*_on_ for nucleosomes than bare DNA substrates^11,17^. In contrast, when bound to solvent-exposed nucleosomal sites, dissociation is ∼10 times slower (*k*_off_) from nucleosomal than free DNA, because GAF-DBD diffusion is confined by core histone-DNA contacts to only ∼5 bp of DNA exposure, and by potential stabilizing histone contacts. Our observation of less frequent but more stable binding to nucleosomal cognate sites is consistent with the dissociation rate compensation mechanism reported for yeast pioneer TFs^17^.

*In vivo*, DNA accessibility is dynamic^57,58^; a particular cognate DNA sequence can alternate between nucleosome-bound and nucleosome-free states owing to the opposing activities of ATP-driven chromatin remodeling enzymes (‘push’ and ‘pull’ from the standpoint of nucleosome linker or NDR DNA)^58,59^. GAF is expressed from an early stage of the *Drosophila* embryo where it is necessary for the establishment and maintenance of nucleosome-depleted sites^29^. Mechanistically, GAF may initially locate cognate sites on a nucleosome core surface by 3D diffusion or a site at the nucleosome edge by 1D invasion (Extended Data Fig. 10a). Upon target recognition, GAF may then recruit remodelers (NURF, PBAP)^60–62^ to translocate nucleosomes away from cognate sites (Extended Data Fig. 10b, c), providing accessibility for subsequent recruitment of the PIC or additional TFs such as HSF at heat shock genes, or Pho and related TFs mediating Polycomb-dependent repression at the corresponding response elements^63^. Whether GAF must dissociate from the nucleosomal target before the remodeler can proceed, or form a ternary complex to prime enzymatic remodeling is unclear. Once cognate site(s) become nucleosome-free, GAF rapidly re-binds by 1D or 3D diffusion and remains on target for minutes^26^. On GAF dissociation, another GAF multimer rapidly substitutes to achieve high occupancy over time, during which the bound TF may act as a physical barrier or roadblock to nucleosome incursions driven by other remodelers (Extended Data Fig. 10d). Persistent occupancy maintains promoter accessibility despite GAF on-off kinetics, enabling downstream PIC recruitment, pausing of RNA Pol II and its release to elongation mode on arrival of additional TFs such as heat shock-activated HSF (Extended Data Fig. 10e, f)^64–69^.

Many DNA and RNA binding proteins search for cognate sites via 1D diffusion, including bacterial TFs^9,59,70–74^ and eukaryotic TF p53^75^. A single GAF-DBD displays rapid back-and-forth, 10 bp-scale 1D diffusion, switching between 2 cognate sites at a rate that is strikingly higher than LacI by 4 orders of magnitude^9^. This indicates that the interaction between a GAF ZF and its cognate sequence is much less stable than LacI, providing an explanation for the requirement for TF multimerization and the natural clustering of multiple GAGAG elements near GAF-dependent promoters and enhancers^37,38^. Our findings highlight 1D sliding at NDRs as a potentially general search mode for other eukaryotic TFs within and beyond the ZF TF family.

In addition, blockage to 1D sliding and the 3D diffusion requirement for robust TF targeting to exposed nucleosome sites has, to our knowledge, hitherto not been demonstrated. Because the bulk of genomic DNA (70-90%) is shielded from 1D search by nucleosome architecture, an optimal balance between 1D and 3D search modes for different TFs will have important implications, especially for TFs that functionally target nucleosome-embedded sites, while an intrinsic aversion to nucleosome association would restrict that TF search space mostly to NDRs. The kinetic interplay between different TF classes, chromatin remodelers and mobilized nucleosomes in dynamically regulating chromatin accessibility provides exciting opportunities for future investigations.

## Acknowledgements

We thank all members of the Ha and Wu laboratories, especially R. Merino Urteaga, J. Hao and T. Liao for support and suggestions, and S. Pangeni, P. Meneses, and C. Carcamo for their optical tweezers experience and offering of many technical suggestions that guided our data collection and analysis. We thank P. Verrijzer for the gift of plasmids. We thank Y. Li at the JHMI Eukaryotic Tissue Culture Facility for FL-GAF expression in SF9 cells. We thank E. Lin and R. He for pilot GAF protein purifications. This study was supported by the National Institutes of Health grants R35 GM149291 (CW), S10 OD025221 (TH) and the Johns Hopkins Discovery Award (CW, GB, TH).

---

**Extended Data Figure 1.**
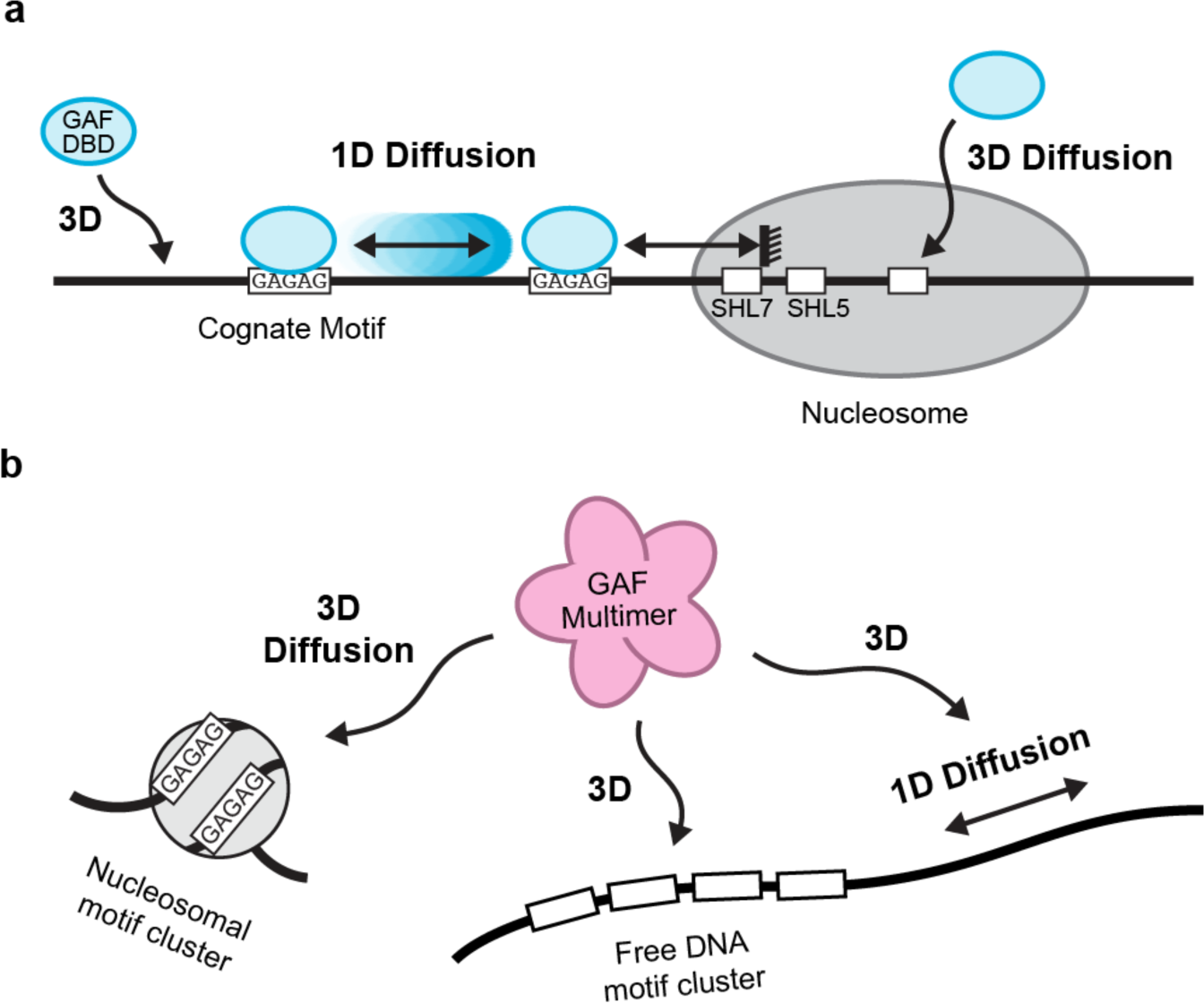
Model for GAF-DBD and GAF-FL target search. **a**, GAF-DBD uses two search modes to locate its target on chromatin. In the 1D sliding mode, GAF-DBD lands on an off-target location on free DNA, then slides back and forth to locate the cognate motif (“GAGAG”). It can escape the cognate site to search for the next site nearby. This 1D search mode allows GAF-DBD to invade into the nucleosome edge but no further. Alternatively, GAF-DBD can also directly associate with a solvent-exposed cognate motif in the nucleosome core from 3D space. This 3D search mode allows GAF to effectively target nucleosomal motifs that are inaccessible by 1D sliding. **b**. GAF-FL uses both 3D and 1D diffusion to locate cognate motif clusters on free DNA. If the motif cluster is inside a nucleosome, GAF-FL can use 3D diffusion for target location.

**Extended Data Figure 2.**
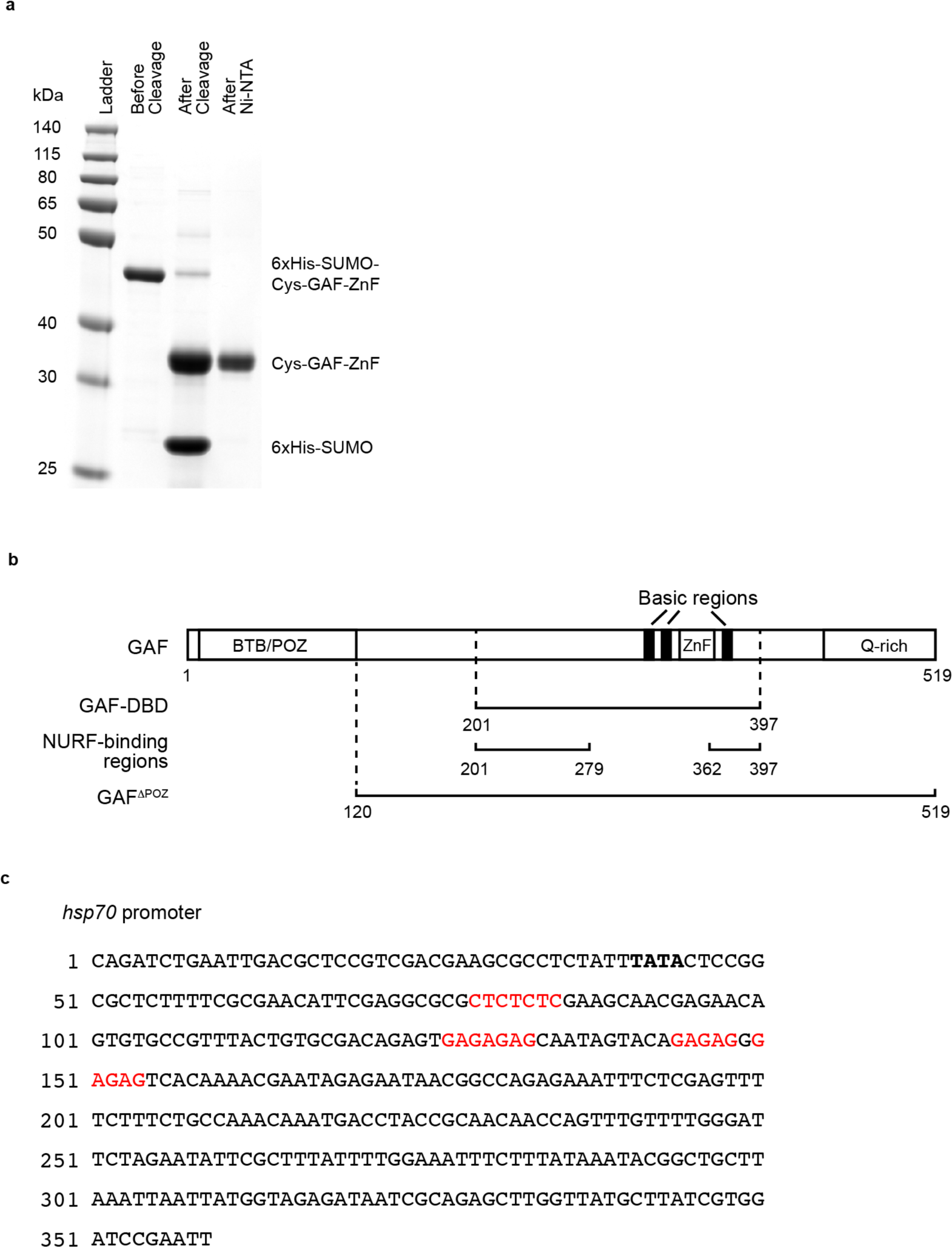
Purification of GAF-DBD protein. **a**, Subtractive Ni-NTA purification after “one-pot” reaction which cleaved off 6xHis-SUMO and labeled the N-terminus of GAF-DBD with Cy3. The calculated molecular weights are 34.4 kDa for 6xHis-SUMO-Cys-GAF-DBD, 21.0 kDa for Cys-GAF-DBD and 13.4 kDa for 6xHis-SUMO. **b**, Schematics of GAF-FL short isoform, GAF-DBD and NURF-binding regions (PMID 11583616) to scale. **c**, Native *hsp70* promoter DNA sequence. GAF cognate sites are highlighted in red. TATA box in bold.

**Extended Data Figure 3.**
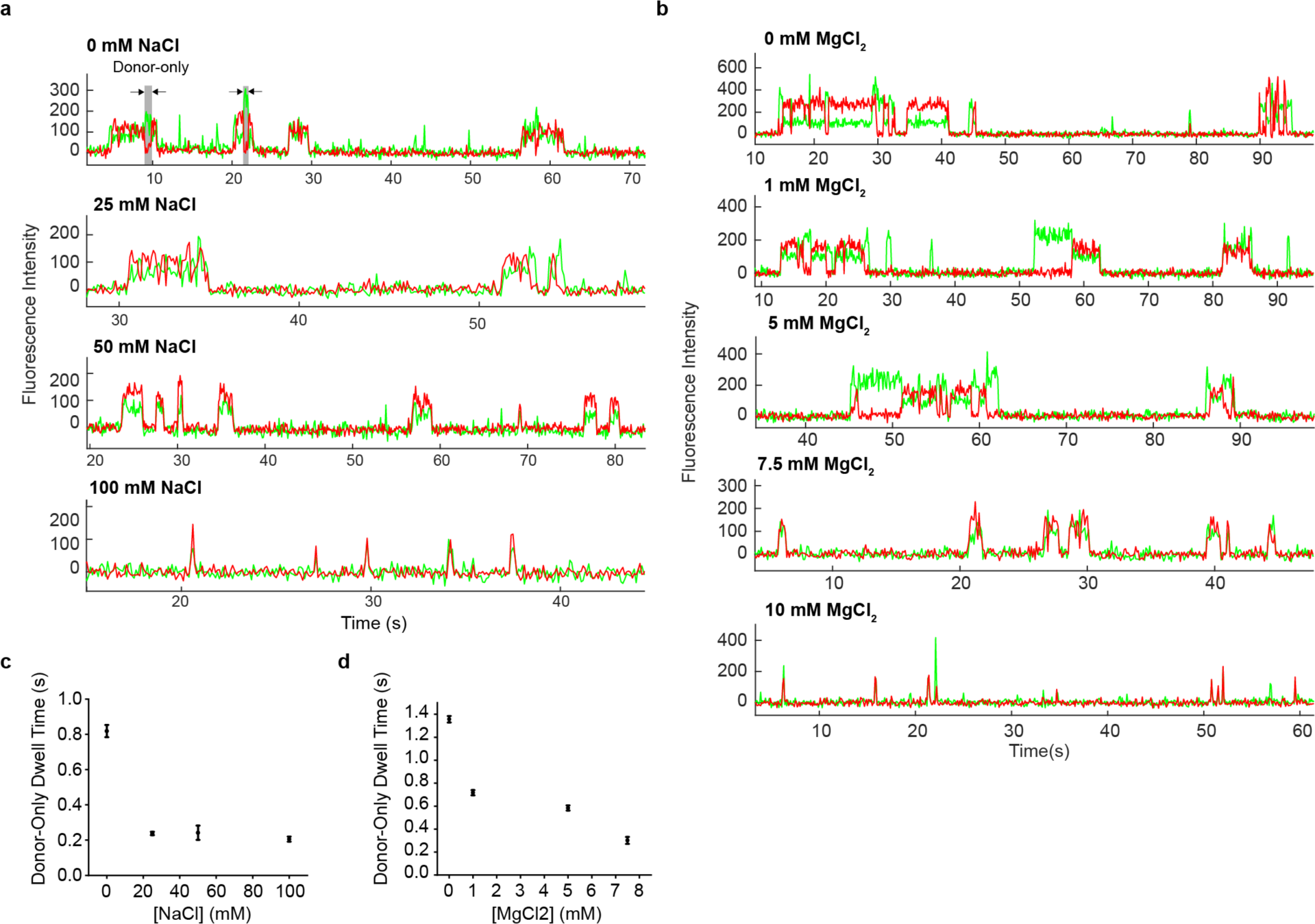
Donor-only dwell time is inversely correlated with ionic strength of buffer solution. **a**, Representative trajectories showing Cy5-cognate DNA bound by Cy3-GAF-DBD in 0, 25, 50 or 100 mM NaCl. Grey-highlighted durations indicate donor-only dwell times. **b**, Representative trajectories showing Cy5-cognate DNA bound by Cy3-GAF-DBD in 0, 1, 5, 7.5, or 10 mM MgCl_2_. **c**, Donor-only dwell time (τ from fitting 1-CDF to a single-exponential decay; see Methods) as a function of NaCl concentration, and **d**, MgCl_2_ concentration. Error bars are standard error from fitting. DNA is the same as cognate DNA in Fig. 1.

**Extended Data Figure 4.**
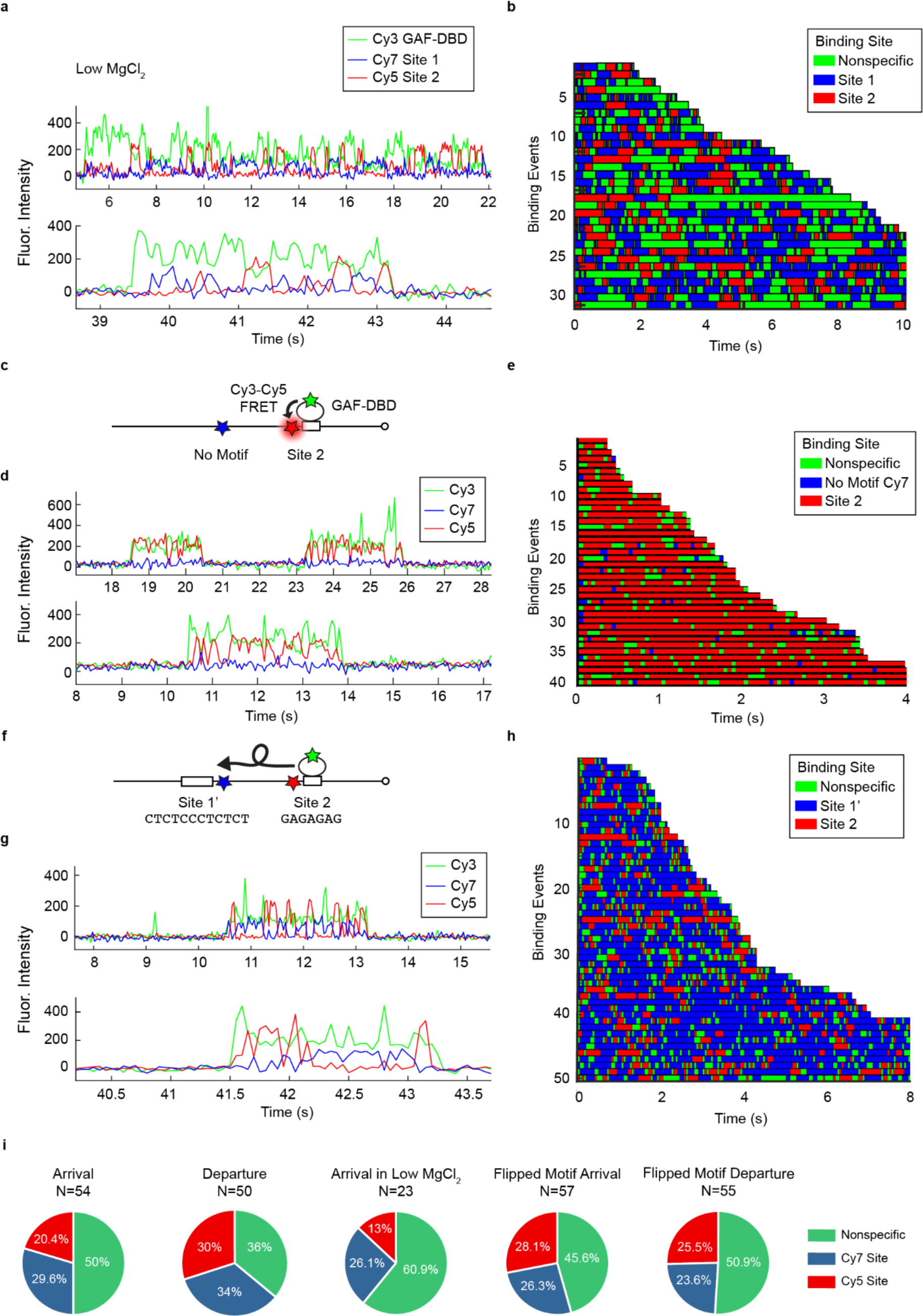
GAF sliding kinetics on Cy5 & Cy7 dual-labeled DNA as a function of salt concentration, cognate site and motif orientation. **a**, Representative single-molecule trajectories of Cy5 & Cy7 DNA bound by Cy3-GAF-DBD in low MgCl_2_ (3 mM). **b**, Rastergram of 31 Cy5 & Cy7 DNA molecules bound by Cy3-GAF-DBD at 3 mM MgCl_2_. **c**, Schematic of Site 2 Only construct where Site 1 was replaced with non-cognate sequence. **d**, Representative single-molecule trajectories of Site 2 Only DNA bound by GAF-DBD. **e**, Rastergram of 40 Site 2 Only DNA molecules bound by GAF-DBD. **f**, Schematic of the flipped motif DNA construct where Site 1 is replaced with its complementary sequence Site 1’. **g**, Representative single-molecule trajectories of flipped motif DNA bound by GAF-DBD. **h**, Rastergram of 57 flipped motif DNA molecules bound by GAF-DBD. DNA same as Fig. 4. **i**, Categories of GAF-DBD arrival landing site and departure launching site.

**Extended Data Figure 5.**
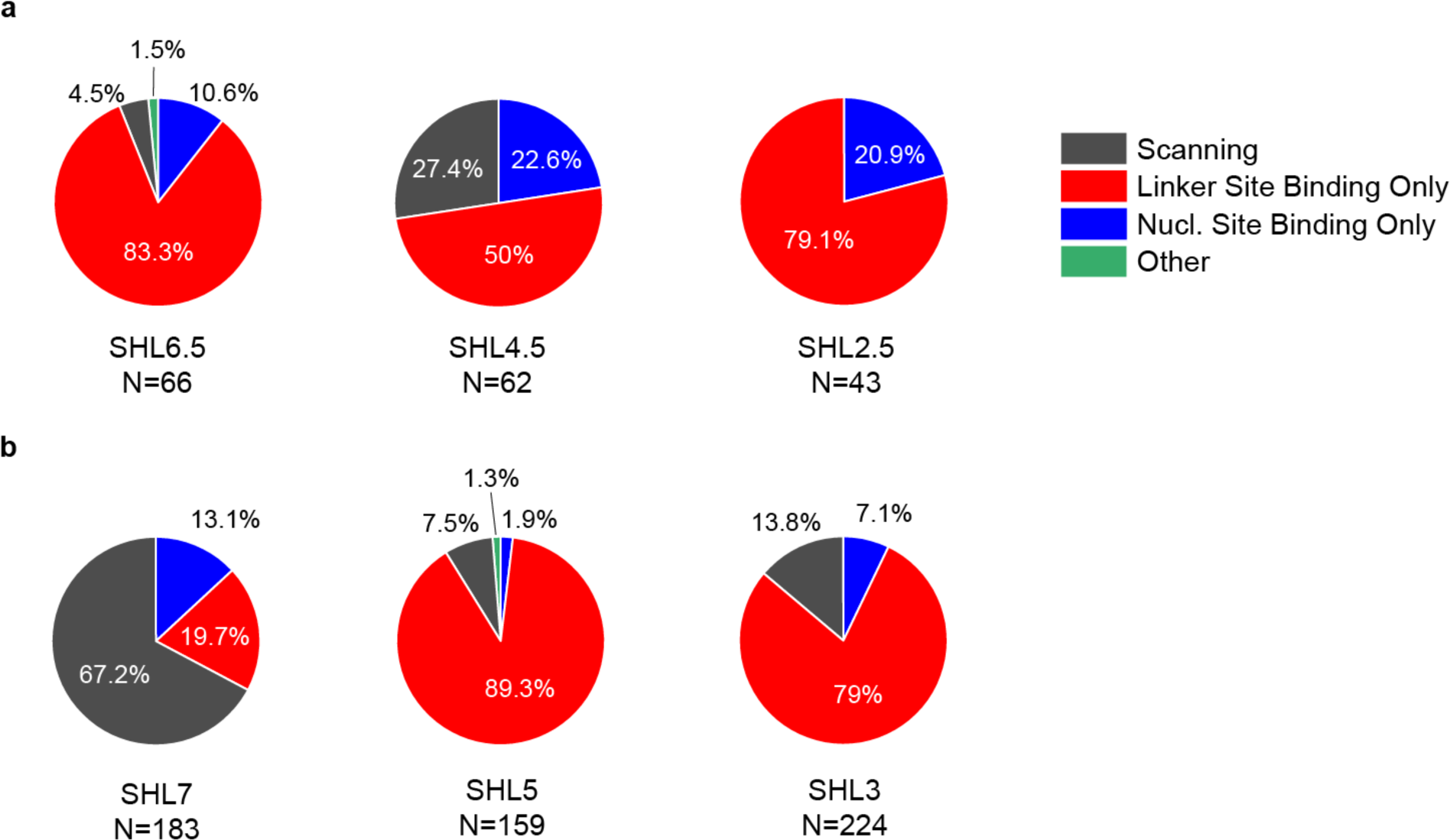
Nucleosomal motif accessibility depends on helical phasing. **a**, Classification of Cy3-GAF-DBD binding events on ‘601’ nucleosomes (40-N-40) with a cognate site placed at SHL6.5, SHL4.5 and SHL2.5. Classification is based on Cy3-Cy7 FRET dynamics, see details in methods. **b**, Classification of Cy3-GAF-DBD binding events on same ‘601’ (40-N-40) nucleosomes with cognate site placed at SHL7, SHL5 and SHL3 (same dataset as Figure 3, re-analyzed for direct comparison with **a**).

**Extended Data Figure 6.**
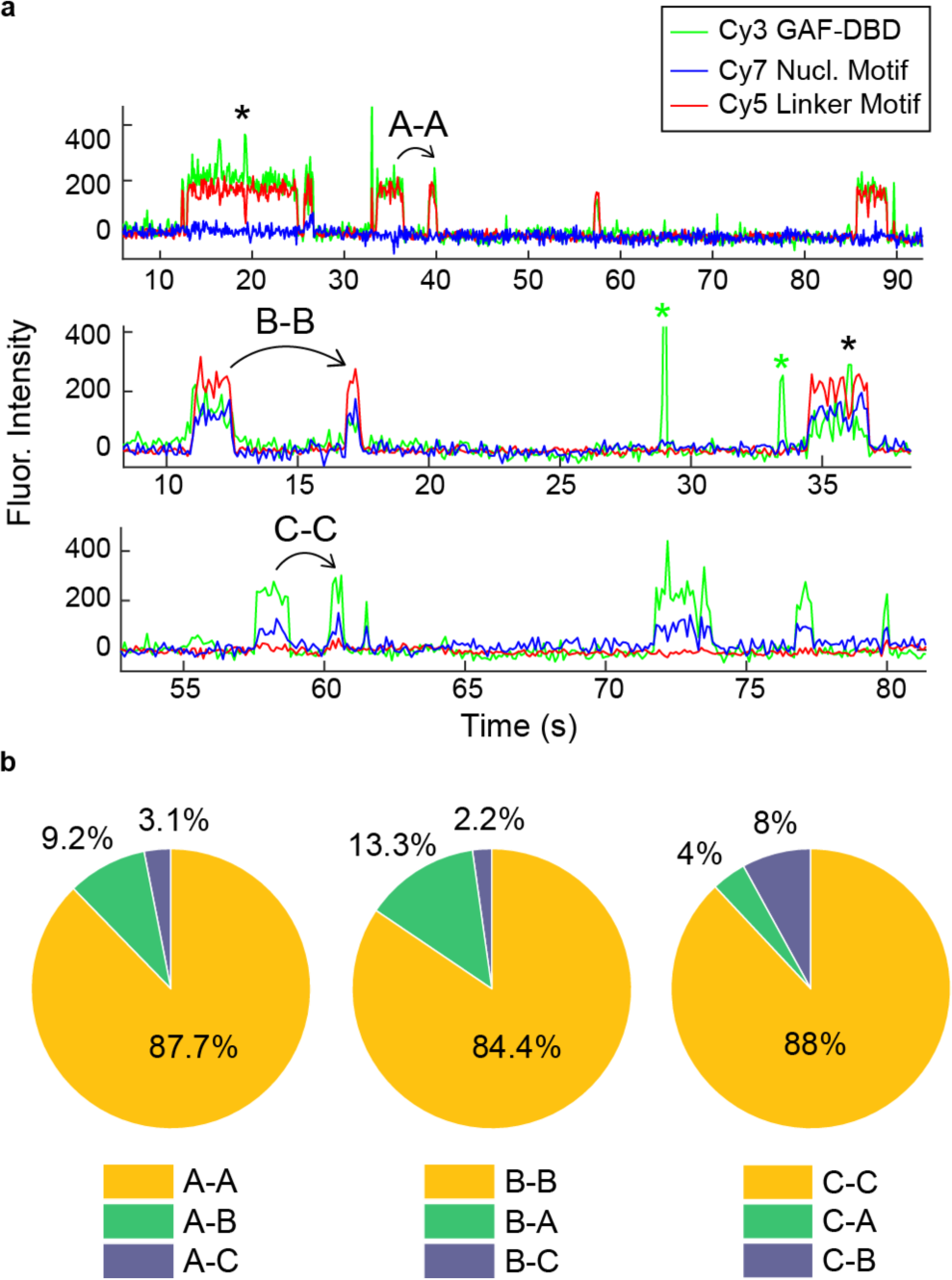
GAF-DBD preferentially re-visits the same cognate site on individual *hsp70* nucleosomes. **a**, Representative single-molecule trajectories showing repetitive visits to the same binding site on a single nucleosome. Upper trace shows repetitive visits to binding site A; middle trace, site B; lower trace, site C. Black asterisks mark transient Cy3 only fluorescence within a binding event, potentially caused by ultra-short-range 1D diffusion on the nucleosome. Green asterisks indicate binding events to non-cognate sites on the nucleosome. **b**, Pie charts showing for all binding events at site A (left pie chart, N = 65), B (middle, N = 45) or C (right, N = 25), the fraction of events that were followed by a second binding to site A, B or C.

**Extended Data Figure 7.**
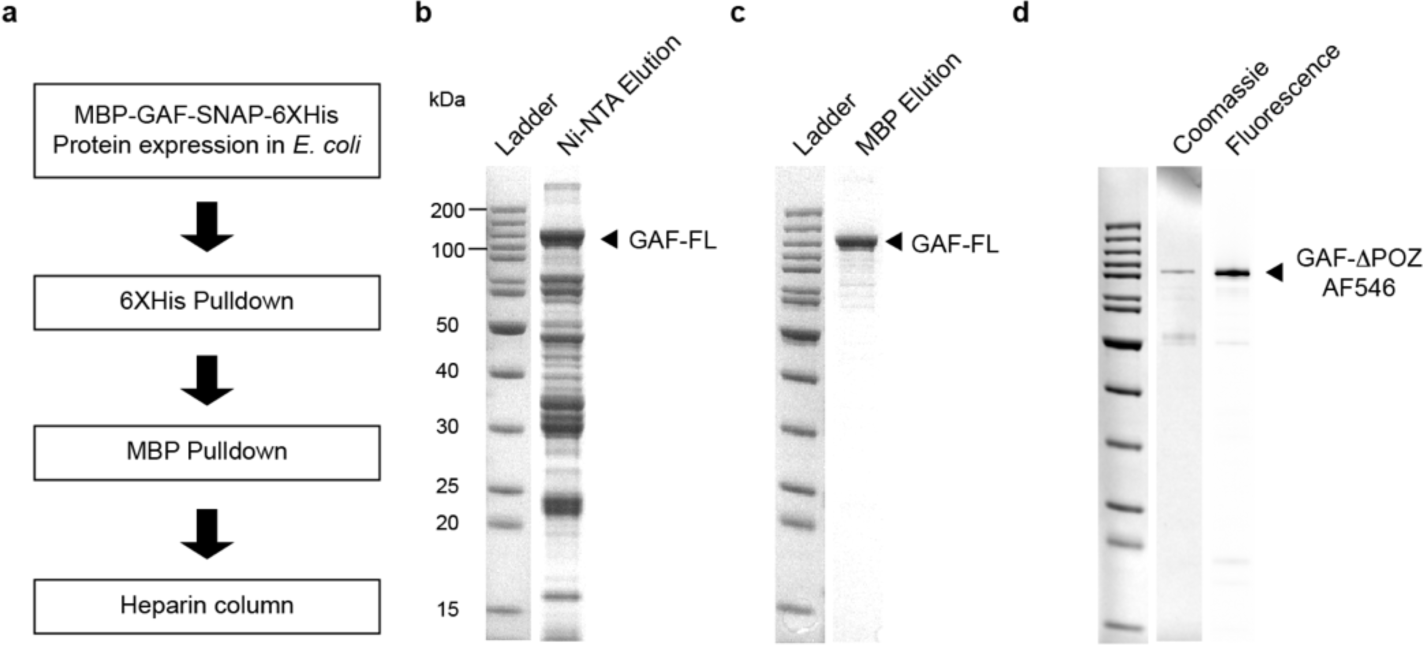
Purification of SNAP-tagged GAF-FL and GAF-ΔPOZ. **a**, GAF-FL purification workflow. **b**, SDS-PAGE gel of GAF-FL elution after 6XHis pulldown, stained with Coomassie Blue. **c**, SDS-PAGE gel of GAF-FL elution after MBP pulldown, stained with Coomassie Blue. **d**, SDS-PAGE gel of AF546-GAF-ΔPOZ, scanned for AlexaFluor 546 fluorescence, then stained with Coomassie Blue.

**Extended Data Figure 8.**
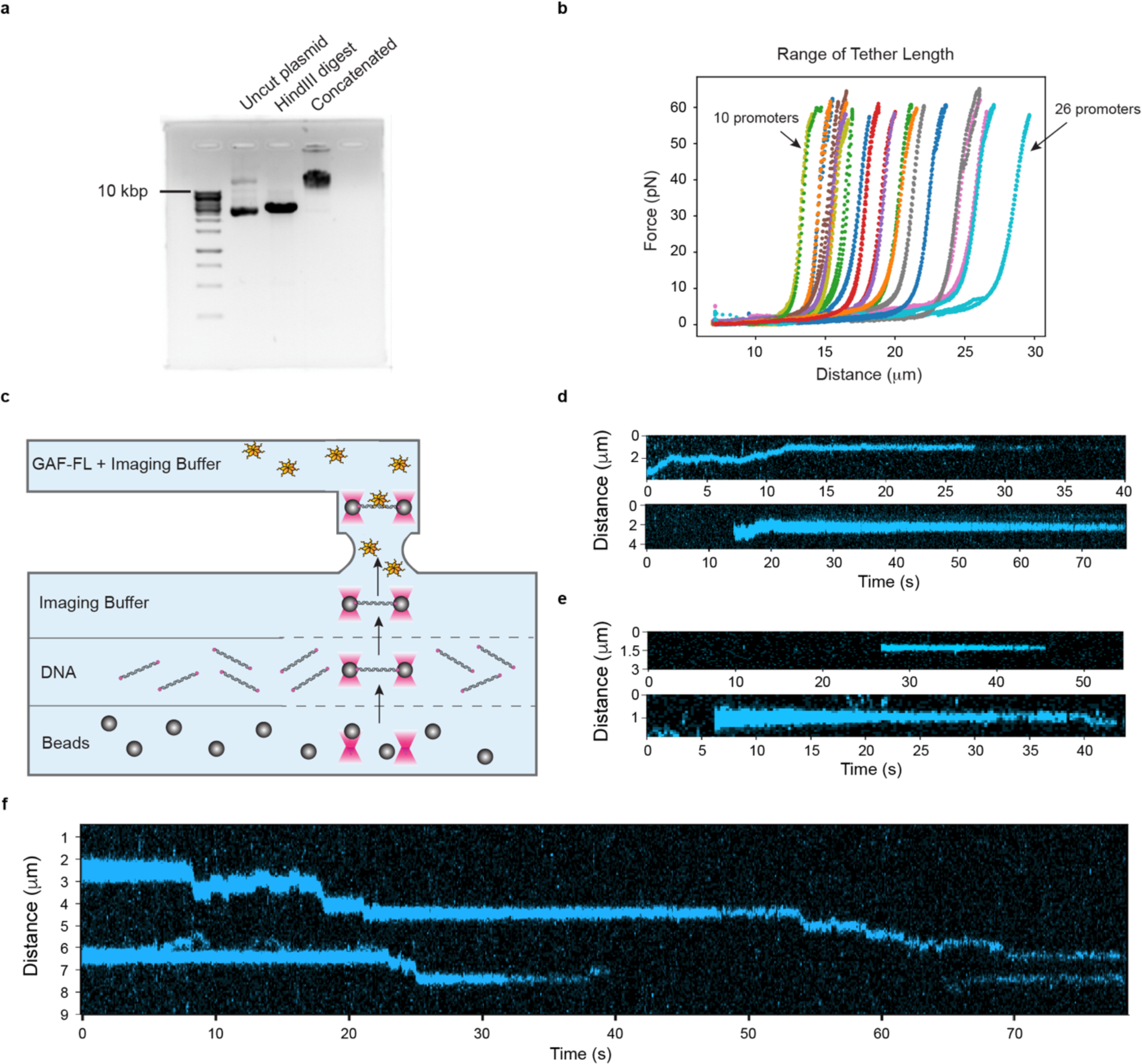
Optical tweezers experiment; design and confirmation of DNA assembly. **a**, Agarose gel electrophoresis of concatenated plasmid DNA. **b**, Force versus distance plot reveals the length of double-stranded DNA tether. **c**, Diagram of flow cell constituents during imaging. Streptavidin coated beads, DNA, and Imaging Buffer were injected to the flow cell under laminar flow. Beads were first optically captured, moved to the DNA channel; once DNA was properly tethered to the trapped beads, the whole assemblage was moved to the protein channel containing AF-488 GAF-FL in Imaging Buffer. Imaging was performed in this channel to maximally visualize binding events. **d**, Representative kymographs where GAF-FL undergoes 1D search on vector+*hsp70* DNA. **e**, Representative kymographs where GAF-FL binds to its target on vector+*hsp70* DNA abruptly from 3D without 1D search. **f**, Representative kymograph where GAF-FL undergoes 1D diffusion from one target to another on vector+*hsp70* DNA.

**Extended Data Figure 9.**
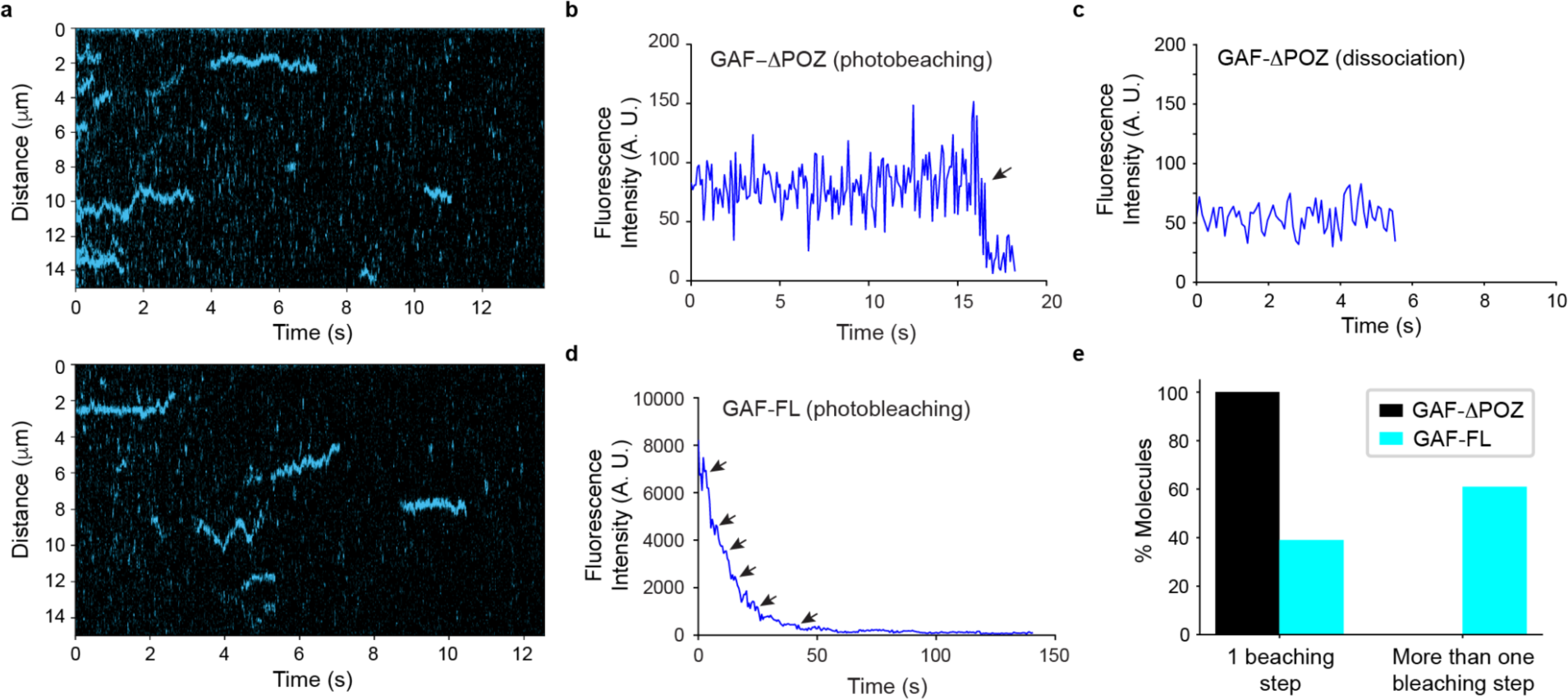
Behavior of full-length GAF depends on multimerization by POZ domain. **a**, Representative kymographs show AF546-GAF-ΔPOZ binding to DNA over time on + *hsp70* DNA. **b**, Representative fluorescence intensity versus time plot for a single trace showing GAF-ΔPOZ photobleaching (indicated by an arrow). **c**, Representative trace showing GAF-ΔPOZ dissociation (at ∼ 5.7 s). Note the abrupt loss of fluorescence signal in this case, which is distinguishable from photobleaching, shown in b, where fluorescence decreases to a near-zero but detectable level. **d**, Representative trace showing GAF-FL photobleaching. Arrows indicate stepwise photobleaching. **e**, Fraction of GAF-ΔPOZ (N=100) and GAF-FL (N=200) molecules with 1 or multiple photobleaching steps.

**Extended Data Figure 10.**
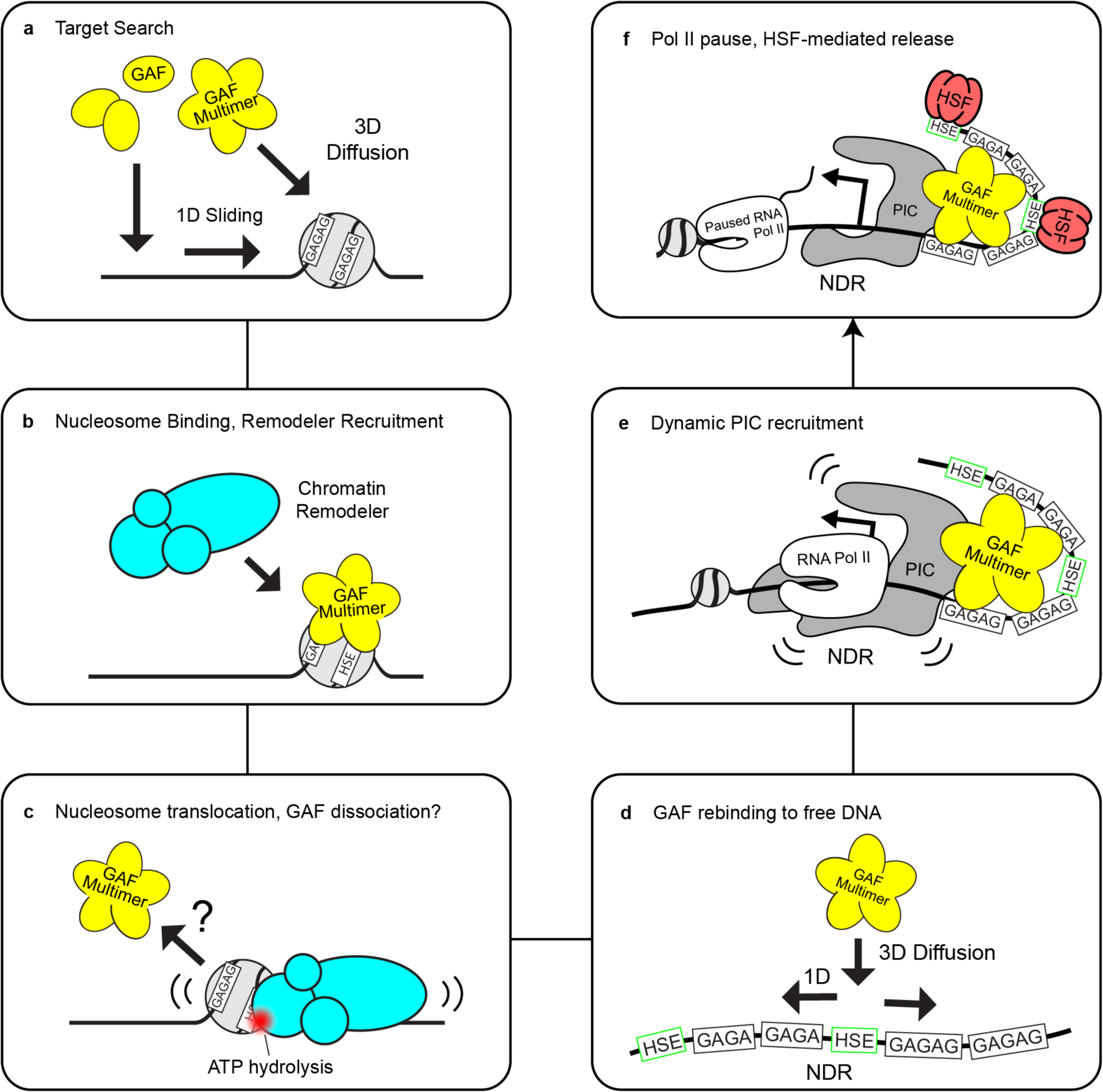
Hypothetical stepwise model for GAF-remodeler collaboration to mobilize targeted nucleosome for PIC assembly.

## Methods

### GAF-DBD protein expression and purification

GAF-DBD (Table 1) was cloned into pET SUMO plasmid (Invitrogen K300-01) using Gibson Assembly (NEB E5520S) to contain an additional N-terminal cysteine for fluorophore labeling. The pET 6xHis-SUMO-Cys-GAF-DBD plasmid was transformed into *E. coli* (Novagen Rosetta 2 DE3, Sigma-Aldrich 71400) for protein expression. On the first day, a starter culture was grown from a single colony in 30 mL Terrific Broth (TB; Sigma-Aldrich T9179) and incubated overnight on a 37°C shaker. One the second day, the starter culture was added to 1 L TB and incubated at 37°C until OD reached 0.8. IPTG was added to 0.4 mM final concentration to induce protein expression. Cell pellets were harvested at 5 hr after IPTG induction and resuspended in 30 mL Lysis Buffer (50 mM Tris pH 7.4, 300 mM NaCl, 10% glycerol, 0.05% Triton X-100), and stored in -80°C.

6xHis-SUMO-Cys-GAF-DBD protein was first purified on Ni-NTA agarose (Qiagen 30210). Briefly, Triton X-100 (0.05% final concentration), protease inhibitor (Roche 4693132001), and β-mercaptoethanol (2 mM final conc.) were added to the thawed cell suspension. Cells were lysed by sonication (amplitude level 50, 10 s on, 50 s off, 3 min process time) on ice. Lysate was clarified by centrifugation at 18,000 g for 45 min at 4°C. The supernatant was transferred to 1.25 mL pre-equilibrated Ni-NTA agarose (2 mL slurry) in a 15 mL tube and incubated at 4°C for 1 hr with gentle shaking. Then agarose beads were washed with 10 mL of Wash Buffer 1 (WB1; 50 mM Tris pH 7.4, 500 mM NaCl, 10% glycerol, 0.05% Triton X-100, 2 mM BME, 20 mM Imidazole pH 8), centrifuged down, resuspended in another 10 mL of WB1, and transferred to a pre-chilled 2.5 cm gravity flow column. Beads were washed three times with 5 mL of WB1, followed by two times with 5 mL of Wash Buffer 2 (50 mM Tris pH 7.4, 300 mM NaCl, 10% glycerol, 0.05% Triton X-100). To elute the protein, 2 mL Elution Buffer (WB2 with 250 mM Imidazole) was added to the beads and incubated for 10 min. 1 mL elution fractions were collected immediately after each addition of elution buffer, then another 1 mL was collected after 5 min incubation. Typically, 10 fractions were collected, and most of the protein eluted in the first 2-3 fractions. Peak fractions were pooled and then further purified by cation exchange chromatography (Cytiva HiTrap SP HP 17115101) on a fast protein liquid chromatography (FPLC) instrument (Cytiva ӒKTA).

### Labeling GAF-DBD with Cy3 fluorophore

We followed the “one-pot” reaction protocol by Jiang et al. (Jiang 2020) to cleave off the 6XHis-SUMO using SUMO protease (Invitrogen K300-01) and label the N-terminal cysteine with Cy3 in a site-specifically manner. Briefly, 6xHis-SUMO-Cys-GAF-DBD was buffer exchanged into Labeling Buffer (100 mM HEPES pH 6.9, 0.5 mM TCEP, 300 mM NaCl) via 24 hr dialysis. In parallel, the transesterification reaction of Cy3-NHS (Cytiva PA13105) was carried out by incubating 0.2 mg of Cy3-NHS in 50 μL of 100 mM HEPES pH 6.9, 0.5 mM TCEP and 500 mM MESNa (Sigma-Aldrich PHR1570) for 6 hr at room temperature, to yield Cy3-MESNa. The “one-pot” labeling and protease cleavage reaction consists of 100 μL of dialyzed protein, 10 μL of SUMO protease, and 12 μL of 6 mM Cy3-MESNa. The reaction proceeded for 36 hr at room temperature. The cleaved 6XHis-SUMO and 6XHis-tagged SUMO protease were removed by Ni-NTA agarose. The labeling efficiency of GAF-DBD was measured to be 92%.

### Insect cell expression and purification of full-length GAF

Expression of FL-GAF in Sf9 cells was performed as described previously (PMID 9927429). Cells harvested from 250 mL Sf9 cell culture were washed twice in ice-cold PBS. For lysis, 150 mL ice-cold lysis buffer (1X PBS, 2mM MgCl_2_ 0.1% Triton X-100, 10% glycerol, 1 mM DTT, 1X protease inhibitor cocktail containing 0.17 μg/mL PMSF, 0.33 μg/mL benzamidine hydrochloride, 1.37 ng/mL pepstatin A, 0.284 ng/mL leupeptin and 2 ng/mL chymostatin) was added to the cell pellet. All further steps were done at 4°C. Cells were resuspended and incubated for 5 min. The nuclei were pelleted by centrifugation at 1,500 g for 4 min. The nuclei pellet was washed once in lysis buffer and salt-extracted in 8 mL lysis buffer supplemented with 300 mM KCl for 1 h. The soluble proteins were collected by centrifugation at 20,000g for 20 min. The nuclear extract was used for anti-HA affinity purification.

A total of 300 μL of 50% anti-hemagglutinin agarose beads (Sigma cat. # A2095) was added to 8 ml of nuclear extract and incubated for 2.5 hr at 4°C with overhead rotation. The beads were washed 5 times with a total of 10 ml of ice-cold lysis buffer. The protein bound to the beads was eluted with 100 μL elution buffer (lysis buffer supplemented with 1 mg/mL HA peptide and additional 850 mM NaCl) for 1 hr at room temperature (RT) with rotation. The final concentration was quantified on a PAGE gel with a bovine serum albumin (BSA) standard curve.

### EMSA of FL-GAF and nucleosome core particle

FL-GAF purified from *Drosophila* SF9 cells was mixed and intubated with 3 nM Cy5-labeled nucleosome core particle (3-N-6-Cy5) reconstituted on hsp70 promoter DNA or Cy5-labeled free DNA at various concentrations in GAF Binding Buffer (12.5 mM HEPES–KOH pH 7.6, 0.05 mM EDTA, 6.25 mM MgCl2, 5% glycerol, 50 mM NaCl, 50 μg/mL BSA (Roche cat. #10711454001), 0.05% NP40) for 1 hr at RT. The samples were added 10% sucrose and loaded onto a 1.3% agarose gel in 0.2X TB (18 mM Tris, 18 mM boric acid) pH 8.0 and run for 45 min at 120 V. The gel was imaged for Cy5 fluorescence on Amersham Typhoon 5 gel imager.

### E. coli expression and purification of SNAP-tagged FL-GAF

MBP-SNAP-GAF-6xHis (SNAP-GAF) was expressed in Lemo21(DE3) Competent *E. coli* (NEB cat. # C2528J). Briefly, the expression plasmid was transformed into Lemo21 cells and a single colony was grown overnight in a 30 mL starter culture, then to 1L in terrific broth. Once

O.D. reaches 0.8, expression was induced by adding IPTG to 400 μM final concentration. The cell pellet was harvested after 5 hr induction at 37°C and resuspended in 30 mL Lysis Buffer (50 mM Tris pH 7.4, 300 mM NaCl, 10% glycerol, 0.05% Triton X-100), and stored in -80°C.

SNAP-GAF protein was purified on Ni-NTA agarose (Qiagen 30210), followed by Amylose resin (NEB cat. # E8021S) and heparin affinity column (Cytiva cat. # 17040601). Because Ni-NTA selects for the C-terminal 6xHis tag and Amiylose selects for the N-terminal MBP tag, this dual-pulldown strategy enriches full-length proteins with intact N- and C-termini. Briefly, Triton X-100 (0.05% final concentration), 1X protease inhibitor (Roche 4693132001), and β-mercaptoethanol (2 mM final conc.) were added to the thawed cell suspension. Cells were lysed by sonication (amplitude level 50, 10 s on, 50 s off, 3 min process time) on ice. Lysate was clarified by centrifugation at 18,000 g for 45 min at 4°C. The supernatant was transferred to 1.25 mL pre-equilibrated Ni-NTA agarose (2 mL slurry) in a 15 mL tube and incubated at 4°C for 1 hr with gentle shaking. Then agarose beads were washed with 10 mL of Wash Buffer 1 (WB1; 50 mM Tris pH 7.4, 500 mM NaCl, 10% glycerol, 0.05% Triton X-100, 2 mM BME, 20 mM Imidazole pH 8), centrifuged down, resuspended in another 10 mL of WB1, and transferred to a pre-chilled 2.5 cm gravity flow column. Beads were washed three times with 5 mL of WB1, followed by two times with 5 mL of Wash Buffer 2 (50 mM Tris pH 7.4, 300 mM NaCl, 10% glycerol, 0.05% Triton X-100). To elute the protein, 2 mL Elution Buffer (WB2 with 250 mM Imidazole) was added to the beads and incubated for 10 min. 1 mL elution fractions were collected immediately after each addition of elution buffer, then another 1 mL was collected after 5 min incubation. Typically, 10 fractions were collected, and most of the protein eluted in the first 2-3 fractions.

To pull down the MBP-tagged SNAP-GAF, peak fractions from the Ni-NTA purification were combined and mixed with pre-equilibrated Amylose resin, then incubated at 4°C for 1 hr with gentle shaking. Amylose resin was then washed with 10 mL of Wash Buffer 1 twice and transferred to a pre-chilled 2.5 cm gravity flow column. The resin was washed 3 times with 5 mL of WB1, followed by 2 times with 5 mL of Wash Buffer 2. To elute the protein, 2 mL MBP Elution Buffer (WB2 + 10 mM maltose) was added and incubated for 10 min. Elution fractions were collected, 1 mL at a time, similar to the Ni-NTA elution step. Peak fractions were pooled and further purified on a heparin column using an FPLC.

### 2-color TIRF microscope

See Poyton & Feng et al. Methods: Two-Color Single-Molecule FRET Microscope Instrumentation.^50^

### 2-color smFRET imaging of Cy3-GAF-DBD binding to Cy5-labeled DNA

Two complementary single-stranded DNA., one containing an internal Cy5 fluorophore, another containing a 5’ biotin, were annealed to form Cy5-labeled DNA constructs. On a homemade PEG-passivated and sparsely biotinylated flow channel (Ref), flow in the following reagents in order: 100 μL of T50 Buffer (10 mM Tris-HCl pH 8, 50 mM NaCl), 40 uL of 0.2 mg/mL Neutravidin in T50 (1 min incubation; Thermo Fisher 31000), 100 uL of T50, 50 μL of 12.5 pM Cy5-DNA (3 min incubation), 100 μL of T50, 50 uL of 0.2 nM Cy3-GAF-DBD in Imaging Buffer (50 mM NaCl, 50 ug/mL BSA (Roche 10711454001), 0.05% NP40 (Sigma I8896), 12.5 mM HEPES–KOH pH 7.6, 0.05 mM EDTA, 6.25 mM MgCl2, 5% glycerol, 0.8% w/v dextrose, 2 mM Trolox, 1 mg/ml glucose oxidase (Sigma-Aldrich G2133) and 500 U/ml catalase (Sigma-Aldrich C3155)). Note that Cy3-GAF-DBD was initially diluted to 20 nM using Storage Buffer (50 mM Tris pH 7.4, 300 mM NaCl, 10% glycerol, 0.05% Triton X-100) to prevent aggregation. The channel was imaged under a 2-color TIRF microscope with 10 Hz frame rate. Laser excitation was programmed to be 10 frames of Cy5 excitation followed by 990 frames of Cy3 excitation for a 100 s movie.

### 2-color smFRET data analysis

2-color movies were converted to single-molecule fluorescence intensity time trajectories using custom-written IDL scripts, and analyzed using custom-written MATLAB scripts. Dwell time data were manually collected by recording the start and end time of each event. The 1-CDF plot of dwell time histogram was fit to a single-exponential decay function (ExpDec1 function: *y* = *y_0_* + *e^−x/τ^*) in Origin. Binding frequency data were manually collected by counting the total number (N) of binding events in each movie and dividing N by the product of movie length and number of trajectories in the movie.

### Labeling DNA oligo with Cy5 or Cy7

Single-stranded DNA oligos were site-specifically labeled with Cy5-NHS (Cytiva PA15100) or Cy7-NHS (Sulfo-Cyanine7-NHS, Lumiprobe 25320) via an amino group attached to an internal thymine through a 6-carbon linker (Integrated DNA Technologies, /iAmMC6T/). A 62.5 μL labeling reaction contains 160 μM amino-modified oligo, 200 mM freshly dissolved NaHCO_3_, 8 mM NHS-dye and nuclease-free water. The reaction was incubated with gentle mixing for 4 hr at room temperature, then overnight at 4°C. The labeled oligo was purified by ethanol precipitation to remove excess dye. Cy7-labeled oligo typically requires two rounds of ethanol precipitation. If the labeling efficiency was lower than 70%, a second round of labeling reaction would be repeated on the labeled oligo. The final labeling efficiencies were typically 80-90%.

### Cy5 Cy7 dual-labeled DNA construction

Making a DNA construct dual-labeled at our desired positions was an engineering challenge because a one-step PCR reaction would require an internally labeled DNA oligo to be longer than 100 bases. Such DNA oligos were not produced by IDT at the time. (Nowadays it can be available by requesting an Ultramer.) To overcome this challenge, a shorter dual-labeled DNA was made first by PCR, restriction digested to produce a sticky overhang, and then ligated with a biotinylated DNA fragment to form the complete construct. Briefly, perform a 2.4 mL PCR reaction using GoTaq Buffer (Promega M7921), 200 μM dNTPs, 1 μM forward primer, 1 μM reverse primer, 100 ng or less template DNA and 24 μL Taq DNA polymerase (NEB M0273L). Purify and concentrate the PCR DNA product by ethanol precipitation. Digest the DNA with DraIII-HF restriction enzyme (NEB R3510L) for 3 hr at 37°C. Purify the digested DNA product by anion exchange chromatography (Cytiva 17115301) on an FPLC instrument, followed by ethanol precipitating DNA in the peak fractions. Simultaneously prepare the biotinylated DNA fragment by annealing two single-stranded DNA oligos. Ligate the Cy5 Cy7 dual-labeled DNA with the biotinylated fragment using T4 DNA ligase (Thermo Fisher EL0011), and purify the ligated DNA by agarose gel extraction.

### Cy5 Cy7 dual-labeled 601 nucleosome reconstitution

See Poyton & Feng et al. Methods: Nucleosome Reconstitution.^50^

### Cy5 Cy7 dual-labeled hsp70p nucleosome reconstitution

The DNA sequence of the hsp70p nucleosome was a modified 187-bp fragment of the *Drosophila melanogaster* hsp70 promoter. According to previously mapped nucleosome positions on a longer hsp70 promoter fragment, our 187 bp DNA should form a 40-N-0 nucleosome, where the 147 bp nucleosomal DNA (N) contains all 9 near-cognate and cognate motifs (GAG, GAGA, GAGAG, etc.) on the 187 bp DNA. To mutate 7 of the motifs, leaving only two GAF motifs intact, we swapped the 7 motifs with the corresponding nucleotides on the Widom 601 sequence (Ref) to preserve nucleosome forming properties of the DNA. To ensure that the nucleosome forms at the desired position on DNA, we followed the nucleosome ligation protocol in (Ref). Briefly, we reconstituted the nucleosome core particle (NCP) using a 150 bp + 3 nt DNA, heat-shifted the NCP at 55°C for 30 min, and then used T4 ligase to attach the 37 bp + 3 nt biotinylated linker DNA by incubating at 4°C for 12 hr.

### 3-color TIRF microscope

See Poyton & Feng et al. Methods: Three-Color Single-Molecule FRET Microscope Instrumentation.^50^

### 3-color imaging of Cy3-GAF-DBD binding to Cy5 & Cy7-labeled DNA or nucleosome

On a homemade PEG-passivated and sparsely biotinylated flow channel (Ref), flow in the following reagents in order: 100 μL of T50 Buffer, 40 uL of 0.2 mg/mL Neutravidin in T50 (1 min incubation), 100 uL of T50, 50 μL of 50 pM Cy5-Cy7-labeled-nucleosome or DNA (3 min incubation), 100 μL of T50, 50 uL of 0.1 nM (unless stated otherwise) Cy3-GAF-DBD in Imaging Buffer (See 2-color imaging methods). The channel was imaged under a 3-color TIRF microscope with 28.6 Hz frame rate (35 ms exposure time per frame) for 601 nucleosomes with GAF motifs (Figure 3) or 10 Hz frame rate for hsp70p nucleosome (Figure 4). Laser excitation was programmed to be 10 frames Cy7, 10 frames Cy5, 960 frames Cy3,10 frames Cy7, 10 frames Cy5.

### 3-color data analysis (dwell time collection, 1-CDF fitting, converting traces to colored stripes)

3-color movies were converted to single-molecule fluorescence intensity time trajectories using custom-written IDL scripts, and analyzed using custom-written MATLAB scripts. Binding event dwell times were manually collected by recording the start and end time of each event. Single-molecule trajectories were converted into rastergrams using custom-written MATLAB scripts for visualization and motif dwell time collection (see next section). The 1-CDF plot of binding event dwell time or motif dwell time histogram was fit to ExpDec1 function in Origin. For Fig. 3, nucleosomes containing with both Cy5 and Cy7 fluorophores were selected for analysis. Binding events were manually categorized into nucleosome and linker DNA binding (FRET alternates between Cy5 and Cy7 in the same binding event), linker DNA binding only (Cy3-Cy5 FRET only), or nucleosome binding only (Cy3-Cy7 FRET only) (Figure 4h). The landing site of GAF-DBD on SHL7-601 (Figure 4i) nucleosome was analyzed by manually categorizing the initial FRET pattern of binding events into 1) landing on linker DNA and sliding to nucleosome (transition from Cy3-Cy5 to Cy3-Cy7 FRET), 2) landing on nucleosome and sliding to linker DNA (transition from Cy3-Cy7 to Cy3-Cy5 FRET), 3) linker DNA binding only (Cy3-Cy5 FRET throughout binding), 4) nucleosome binding only (Cy3-Cy7 FRET throughout binding), 5) nonspecific binding and sliding to linker DNA (Cy3 signal only to Cy3-Cy5 FRET), and 6) nonspecific binding and sliding to nucleosome (Cy3 signal only to Cy3-Cy7 FRET).

For Extended Data Fig. 5, nucleosomes with Cy7 fluorophore were selected for analysis. Binding events were categorized into 1) linker motif binding only if it shows constant Cy3-Cy5 FRET and no Cy3-Cy7 FRET throughout, 2) nucleosomal motif binding only if it shows constant Cy3-Cy7 FRET and no Cy3-Cy5 FRET throughout, 3) scanning if it exhibits dynamic transitions between Cy3-Cy7 FRET and either Cy3 only or Cy3-Cy5 FRET during binding, or 4) other if none of the above applies.

### Binding site assignment for 3-color FRET data

3-color FRET trajectories with clear alternating Cy5/Cy7 emission (Fig. 2, Fig. 3 SHL7, Extended Data Fig. 4b, 4h) were fit to a hidden markov model (see Supplementary Note) for binding site assignments. The rest of the trajectories (Fig. 3 SHL5 and SHL3, Extended Data Fig. 4e) were analyzed with a custom-written Relative Intensity Algorithm. Briefly, the algorithm assigns binding at a specific time point to Cy5-site if Cy5 fluorescence is stronger than Cy7, or to Cy7-site if Cy7 fluorescence is stronger. If it detects only Cy3 fluorescence, then nonspecific binding is assigned.

### Biotinylated concatenated plasmid DNA preparation

The Drosophila hsp70 promoter sequence was inserted into the 3.32 pBluescript cloning vector. 25μg of the plasmid containing the hsp70 promoter sequence was digested with HindIII restriction enzyme (NEB R3104) for 3 hours at 37C. The digested fragment was then purified using a Qiagen DNA purification kit (QIAGEN-20021). 6μg the purified fragments were then concatenated and biotinylated using the “Adaptor 1” Lumicks DNA repeat assembly kit designed for HindIII overhangs (Lumicks 00029). The same was performed for the empty vector control that does not contain the hsp70 promoter sequence or any other GAGAG motifs. Success of assembly reaction was confirmed by gel shift on agarose gel (Extended Data Fig. X1a).

### Optical Tweezers and Confocal Microscopy Data Collection

Optical tweezers experiments were performed at room temperature on a LUMICKS C-Trap configured with two optical traps. Imaging was performed using a LUMICKS 5-channel microfluidic flow cell as previously described (54,1). Due to the heterogeneity in concatenated plasmid length, DNA tethered between the beads was first checked by force versus distance curve to ensure a proper double stranded DNA and identify the distance between the beads. FD Curves were collected by bringing the beads 5μm apart, zeroing the force, and extended to maximum stretch at 60pN and exported as FD Curves from the bluelake software and plotted using Lumicks pylake (Extended Data Fig. 8b). Distance between the beads at a force of 5pN was used for determining the length of the DNA tether and number of hsp70 promoters present (Extended Data Fig. 8b). Traps were then moved to the protein channel and protein was flown on the tether stretched to 5pN at 0.3 bar. AF488-GAF-FL or AF546-GAF-ΔPOZ was flown through the protein channel diluted to 80 nM in imaging buffer (50 mM NaCl, 50g/mL BSA (Roche 10711454001), 0.05% NP40 (Sigma I8896), 12.5 mM HEPES-KOH pH 7.6, 0.5 mM EDTA, 6.25 mM MgCl2, 0.8% w/v dextrose, and 2 mM Trolox). Flow was turned off before imaging. Kymographs were generated by setting a confocal scan line between the beads at a pixel time of 0.1 ms and size of 100 nm. Due to the differences in the DNA tether length from heterogenous concatenation, the confocal scan line was modified before imaging each new tether to make sure it spans the entire length of the DNA. This resulted in a frame rate range of 70-200 ms.

### Single Particle Tracking and Diffusion Analysis

Raw kymograph files were processed using the Lumicks custom software Pylake v.1.0.0 python package. All scripts for kymotracking and diffusion analysis are written as detailed in the Lumicks Pylake user guide (https://lumicks-pylake.readthedocs.io/en/v1.0.0/theory/diffusion/ diffusion.html). Molecules were tracked using the kymotrack widget with centroid refinement and track width set to 0.5. Output of each tracked molecule gave position, time, and summed photon count that were used for downstream analysis. MSD values were obtained for each trace using the Lumicks “msd” function with maximum lags set to 5. Because molecules exhibited the same diffusion pattern in each condition, averages of the MSD values for each time lag from each trace were reported (Fig. 5g). Diffusion constant was measured using the Lumicks “estimate_diffusion” function for ordinary least squares linear regression of the MSD values. This was performed for each trace and the diffusion constant was averaged for each condition. Statistical analysis was performed using an unpaired Student’s t-test in prism, p-values are reported in figure legends.

## Notes

### Competing Interest Statement

The authors have declared no competing interest.

### Summary of Updates

Substantial update on data, figures, and text.

## References

1. Flury, V. & Groth, A. Safeguarding the epigenome through the cell cycle: a multitasking game. Curr. Opin. Genet. Dev. 85, 102161 (2024).

2. Li, B., Carey, M. & Workman, J. L. The role of chromatin during transcription. Cell 128, 707–719 (2007).

3. Kadonaga, J. T. Eukaryotic transcription: an interlaced network of transcription factors and chromatin-modifying machines. Cell 92, 307–313 (1998).

4. Isbel, L., Grand, R. S. & Schübeler, D. Generating specificity in genome regulation through transcription factor sensitivity to chromatin. Nat. Rev. Genet. 23, 728–740 (2022).

5. Lukas, J., Lukas, C. & Bartek, J. More than just a focus: The chromatin response to DNA damage and its role in genome integrity maintenance. Nat. Cell Biol. 13, 1161–1169 (2011).

6. von Hippel, P. H. & Berg, O. G. Facilitated target location in biological systems. J. Biol. Chem. 264, 675–678 (1989).

7. Slutsky, M. & Mirny, L. A. Kinetics of protein-DNA interaction: facilitated target location in sequence-dependent potential. Biophys. J. 87, 4021–4035 (2004).

8. Hettich, J. & Gebhardt, J. C. M. Transcription factor target site search and gene regulation in a background of unspecific binding sites. J. Theor. Biol. 454, 91–101 (2018).

9. Marklund, E. et al. DNA surface exploration and operator bypassing during target search. Nature 583, 858–861 (2020).

10. Suter, D. M. Transcription Factors and DNA Play Hide and Seek. Trends Cell Biol. 30, 491– 500 (2020).

11. Luo, Y., North, J. A., Rose, S. D. & Poirier, M. G. Nucleosomes accelerate transcription factor dissociation. Nucleic Acids Res. 42, 3017–3027 (2014).

12. Zhu, F. et al. The interaction landscape between transcription factors and the nucleosome. Nature 562, 76–81 (2018).

13. Michael, A. K. et al. Mechanisms of OCT4-SOX2 motif readout on nucleosomes. Science 368, 1460–1465 (2020).

14. Michael, A. K. et al. Cooperation between bHLH transcription factors and histones for DNA access. Nature 619, 385–393 (2023).

15. Li, S., Zheng, E. B., Zhao, L. & Liu, S. Nonreciprocal and Conditional Cooperativity Directs the Pioneer Activity of Pluripotency Transcription Factors. Cell Rep. 28, 2689–2703.e4 (2019).

16. Donovan, B. T. et al. Basic helix-loop-helix pioneer factors interact with the histone octamer to invade nucleosomes and generate nucleosome-depleted regions. Mol. Cell 83, 1251– 1263.e6 (2023).

17. Donovan, B. T., Chen, H., Jipa, C., Bai, L. & Poirier, M. G. Dissociation rate compensation mechanism for budding yeast pioneer transcription factors. Elife 8, (2019).

18. Mivelaz, M. et al. Chromatin Fiber Invasion and Nucleosome Displacement by the Rap1 Transcription Factor. Mol. Cell 77, 488–500.e9 (2020).

19. Biggin, M. D. & Tjian, R. Transcription factors that activate the Ultrabithorax promoter in developmentally staged extracts. Cell 53, 699–711 (1988).

20. Gilmour, D. S., Thomas, C. & Elgin, S. C. Drosophila nuclear proteins bind to regions of alternating C and T residues in gene promoters. Science 245, 1487–1490 (1989).

21. Farkas, G. et al. The Trithorax-like gene encodes the Drosophila GAGA factor. Nature 371, 806–808 (1994).

22. Soeller, W. C., Oh, C. E. & Kornberg, T. B. Isolation of cDNAs encoding the Drosophila GAGA transcription factor. Mol. Cell. Biol. 13, 7961–7970 (1993).

23. Tsukiyama, T., Becker, P. B. & Wu, C. ATP-dependent nucleosome disruption at a heat-shock promoter mediated by binding of GAGA transcription factor. Nature 367, 525–532 (1994).

24. Omichinski, J. G., Pedone, P. V., Felsenfeld, G., Gronenborn, A. M. & Clore, G. M. The solution structure of a specific GAGA factor-DNA complex reveals a modular binding mode. Nat. Struct. Biol. 4, 122–132 (1997).

25. Fuda, N. J. et al. GAGA factor maintains nucleosome-free regions and has a role in RNA polymerase II recruitment to promoters. PLoS Genet. 11, e1005108 (2015).

26. Tang, X. et al. Kinetic principles underlying pioneer function of GAGA transcription factor in live cells. Nat. Struct. Mol. Biol. 29, 665–676 (2022).

27. Adkins, N. L., Hagerman, T. A. & Georgel, P. GAGA protein: a multi-faceted transcription factor. Biochem. Cell Biol. 84, 559–567 (2006).

28. Tsukiyama, T. & Wu, C. Purification and properties of an ATP-dependent nucleosome remodeling factor. Cell 83, 1011–1020 (1995).

29. Gaskill, M. M., Gibson, T. J., Larson, E. D. & Harrison, M. M. GAF is essential for zygotic genome activation and chromatin accessibility in the early embryo. Elife 10, (2021).

30. Chetverina, D., Erokhin, M. & Schedl, P. GAGA factor: a multifunctional pioneering chromatin protein. Cell. Mol. Life Sci. 78, 4125–4141 (2021).

31. Wilkins, R. C. & Lis, J. T. DNA distortion and multimerization: novel functions of the glutamine-rich domain of GAGA factor. J. Mol. Biol. 285, 515–525 (1999).

32. Mahmoudi, T., Katsani, K. R. & Verrijzer, C. P. GAGA can mediate enhancer function in trans by linking two separate DNA molecules. EMBO J. 21, 1775–1781 (2002).

33. Li, X. et al. GAGA-associated factor fosters loop formation in the Drosophila genome. Mol. Cell 83, 1519–1526.e4 (2023).

34. Duarte, F. M. et al. Transcription factors GAF and HSF act at distinct regulatory steps to modulate stress-induced gene activation. Genes Dev. 30, 1731–1746 (2016).

35. Batut, P. J. et al. Genome organization controls transcriptional dynamics during development. Science 375, 566–570 (2022).

36. Lambert, S. A. et al. The Human Transcription Factors. Cell 172, 650–665 (2018).

37. Katsani, K. R., Hajibagheri, M. A. & Verrijzer, C. P. Co-operative DNA binding by GAGA transcription factor requires the conserved BTB/POZ domain and reorganizes promoter topology. EMBO J. 18, 698–708 (1999).

38. van Steensel, B., Delrow, J. & Bussemaker, H. J. Genomewide analysis of Drosophila GAGA factor target genes reveals context-dependent DNA binding. Proc. Natl. Acad. Sci. U. S. A. 100, 2580–2585 (2003).

39. Ha, T. et al. Probing the interaction between two single molecules: fluorescence resonance energy transfer between a single donor and a single acceptor. Proc. Natl. Acad. Sci. U. S. A. 93, 6264–6268 (1996).

40. Ha, T. et al. Fluorescence resonance energy transfer at the single-molecule level. Nature Reviews Methods Primers 4, 1–18 (2024).

41. Jiang, H., D’Agostino, G. D., Cole, P. A. & Dempsey, D. R. Selective protein N-terminal labeling with N-hydroxysuccinimide esters. Methods Enzymol. 639, 333–353 (2020).

42. Bonnet, I. et al. Sliding and jumping of single EcoRV restriction enzymes on non-cognate DNA. Nucleic Acids Res. 36, 4118–4127 (2008).

43. Moll, J. R., Acharya, A., Gal, J., Mir, A. A. & Vinson, C. Magnesium is required for specific DNA binding of the CREB B-ZIP domain. Nucleic Acids Res. 30, 1240–1246 (2002).

44. Lowary, P. T. & Widom, J. New DNA sequence rules for high affinity binding to histone octamer and sequence-directed nucleosome positioning. J. Mol. Biol. 276, 19–42 (1998).

45. Li, G., Levitus, M., Bustamante, C. & Widom, J. Rapid spontaneous accessibility of nucleosomal DNA. Nat. Struct. Mol. Biol. 12, 46–53 (2005).

46. Huh, J.-W. et al. Multivalent di-nucleosome recognition enables the Rpd3S histone deacetylase complex to tolerate decreased H3K36 methylation levels. EMBO J. 31, 3564– 3574 (2012).

47. Hamiche, A., Sandaltzopoulos, R., Gdula, D. A. & Wu, C. ATP-dependent histone octamer sliding mediated by the chromatin remodeling complex NURF. Cell 97, 833–842 (1999).

48. Georgel, P. T. Chromatin potentiation of the hsp70 promoter is linked to GAGA-factor recruitment. Biochem. Cell Biol. 83, 555–565 (2005).

49. Sabantsev, A., Levendosky, R. F., Zhuang, X., Bowman, G. D. & Deindl, S. Direct observation of coordinated DNA movements on the nucleosome during chromatin remodelling. Nat. Commun. 10, 1720 (2019).

50. Poyton, M. F. et al. Coordinated DNA and histone dynamics drive accurate histone H2A.Z exchange. Sci Adv 8, eabj5509 (2022).

51. Stinson, B. M., Moreno, A. T., Walter, J. C. & Loparo, J. J. A Mechanism to Minimize Errors during Non-homologous End Joining. Mol. Cell 77, 1080–1091.e8 (2020).

52. Polach, K. J. & Widom, J. Mechanism of protein access to specific DNA sequences in chromatin: a dynamic equilibrium model for gene regulation. J. Mol. Biol. 254, 130–149 (1995).

53. Li, G. & Widom, J. Nucleosomes facilitate their own invasion. Nat. Struct. Mol. Biol. 11, 763–769 (2004).

54. Rudnizky, S., Khamis, H., Malik, O., Melamed, P. & Kaplan, A. The base pair-scale diffusion of nucleosomes modulates binding of transcription factors. Proc. Natl. Acad. Sci. U. S. A. 116, 12161–12166 (2019).

55. Tan, C. & Takada, S. Nucleosome allostery in pioneer transcription factor binding. Proc. Natl. Acad. Sci. U. S. A. 117, 20586–20596 (2020).

56. Judd, J., Duarte, F. M. & Lis, J. T. Pioneer-like factor GAF cooperates with PBAP (SWI/SNF) and NURF (ISWI) to regulate transcription. Genes Dev. 35, 147–156 (2021).

57. Calderon, D. et al. The continuum of embryonic development at single-cell resolution. Science 377, eabn5800 (2022).

58. Kubik, S. et al. Opposing chromatin remodelers control transcription initiation frequency and start site selection. Nat. Struct. Mol. Biol. 26, 744–754 (2019).

59. Kim, J. M. et al. Dynamic 1D search and processive nucleosome translocations by RSC and ISW2 chromatin remodelers. Elife 12, (2024).

60. Okada, M. & Hirose, S. Chromatin remodeling mediated by Drosophila GAGA factor and ISWI activates fushi tarazu gene transcription in vitro. Mol. Cell. Biol. 18, 2455–2461 (1998).

61. Nakayama, T., Shimojima, T. & Hirose, S. The PBAP remodeling complex is required for histone H3.3 replacement at chromatin boundaries and for boundary functions. Development 139, 4582–4590 (2012).

62. Xiao, H. et al. Dual functions of largest NURF subunit NURF301 in nucleosome sliding and transcription factor interactions. Mol. Cell 8, 531–543 (2001).

63. Brown, J. L., Price, J. D., Erokhin, M. & Kassis, J. A. Context-dependent role of Pho binding sites in Polycomb complex recruitment in Drosophila. Genetics 224, (2023).

64. Badenhorst, P., Voas, M., Rebay, I. & Wu, C. Biological functions of the ISWI chromatin remodeling complex NURF. Genes Dev. 16, 3186–3198 (2002).

65. Boija, A. et al. CBP Regulates Recruitment and Release of Promoter-Proximal RNA Polymerase II. Mol. Cell 68, 491–503.e5 (2017).

66. Kuroda, M. I., Kang, H., De, S. & Kassis, J. A. Dynamic Competition of Polycomb and Trithorax in Transcriptional Programming. Annu. Rev. Biochem. 89, 235–253 (2020).

67. Wolle, D. et al. Functional Requirements for Fab-7 Boundary Activity in the Bithorax Complex. Mol. Cell. Biol. 35, 3739–3752 (2015).

68. Brahma, S. & Henikoff, S. RNA Polymerase II, the BAF remodeler and transcription factors synergize to evict nucleosomes. bioRxiv (2023) doi:10.1101/2023.01.22.525083.

69. Nguyen, V. Q. et al. Spatiotemporal coordination of transcription preinitiation complex assembly in live cells. Mol. Cell 81, 3560–3575.e6 (2021).

70. Carcamo, C. C. et al. ATP binding facilitates target search of SWR1 chromatin remodeler by promoting one-dimensional diffusion on DNA. Elife 11, (2022).

71. Gowers, D. M., Wilson, G. G. & Halford, S. E. Measurement of the contributions of 1D and 3D pathways to the translocation of a protein along DNA. Proc. Natl. Acad. Sci. U. S. A. 102, 15883–15888 (2005).

72. Ragunathan, K., Liu, C. & Ha, T. RecA filament sliding on DNA facilitates homology search. Elife 1, e00067 (2012).

73. Globyte, V., Lee, S. H., Bae, T., Kim, J.-S. & Joo, C. CRISPR/Cas9 searches for a protospacer adjacent motif by lateral diffusion. EMBO J. 38, (2019).

74. Chandradoss, S. D., Schirle, N. T., Szczepaniak, M., MacRae, I. J. & Joo, C. A Dynamic Search Process Underlies MicroRNA Targeting. Cell 162, 96–107 (2015).

75. Tafvizi, A. et al. Tumor suppressor p53 slides on DNA with low friction and high stability. Biophys. J. 95, L01–3 (2008).

