## Supplemental Information for "GAGA zinc finger transcription factor searches chromatin by 1D-3D facilitated diffusion"

### 1. **GAF-ZnF Protein Sequence** (**C** indicates labeling position)

CVIQAFLPARKRKPRVKKMSPTAPKISKVEGMDTIMGTPTSSHGSGSVQQVLGENGAEGQLLSS
TPIIKSEGQKVETIVTMDPNNMIPVTSANAATGEITPAQGATGSSGGNTSGVLSTPKAKRAKHP
PGTEKPRSRSQSEQPATCPICYAVIRQSRNLRRHLELRHFAKPGVKKEKKSKSGNDTTLDSSME
MNTTAE

### 2. **Cy5-DNA Oligos (**GAF cognate motif is highlighted)

| | | |
|---|---|---|
| Cognate-Motif | Strand 1 | /5Biosg/GT TTT GTG AGT ACG CGC TGT TGT ACT ATT GTT AAC CCA CAA GGT CGC ACA GTA AAC GGC ACA CTG TAC GTG TTG CTT CGA GAG AGC GCG C |
| | Strand 2 | GCG CGC TCT CTC GAA GC/iCy5/ AAC ACG TAC AGT GTG CCG TTT ACT GTG CGA CCT TGT GGG TTA ACA ATA GTA CAA CAG CGC GTA CTC ACA AAA C |
| Non-Motif | Strand 1 | /5Biosg/GT TTT GTG AGT ACG CGC TGT TGT ACT ATT GTT AAC CCA CAA GGT CGC ACA GTA AAC GGC ACA CTG TAC GTG TTG CTT CTA TAC AGC GCG C |
| | Strand 2 | GCG CGC TGT ATA GAA GC/iCy5/ AAC ACG TAC AGT GTG CCG TTT ACT GTG CGA CCT TGT GGG TTA ACA ATA GTA CAA CAG CGC GTA CTC ACA AAA C |

Table 1. Cy5-DNA oligos for two-color FRET.

### 3. **Cy5 & Cy7 Dual-Labeled DNA Constructs**

*601 DNA Constructs and Primers*

Annotations:

| | |
|---|---|
| **t** | Cy5 labeling position |
| **A** | Cy7 labeling position |
| GAGAGA | GAF cognate motif |
| CACCTGGTG | DraIII restriction site |
| lowercase | Linker DNA |
| UPPERCASE | Nucleosomal DNA |

| | |
|---|---|
| 601-SHL7 Final | tatcc**t**actggagagagcaaggtcgctgttcaatacatgcAGAGAGAGT**A**TA TATCTGACACGTGCCTGGAGACTAGGGAGTAATCCCCTTGGCGGTTACACCT GGTGGGACAGCGCGTACGTGCGTTTAAGCGGTGCTAGAGCTGTCTACGACCA ATTGAGCGGCCTCGGCACCGGGATTCTCCAGggcggccgcgtatagggtcca tcacataagggatgaactc |
| 601-SHL5 Final | tatcc**t**actggagagagcaaggtcgctgttcaatacatgcACAGGATGTATA TATCTGACAGAGAGAT**A**GAGACTAGGGAGTAATCCCCTTGGCGGTTACACCT GGTGGGACAGCGCGTACGTGCGTTTAAGCGGTGCTAGAGCTGTCTACGACCA |

| | |
|---|---|
| | ATTGAGCGGCCTCGGCACCGGGATTCTCCAGggcggccgcgtatagggtcca<br>tcacataagggatgaactc |
| 601-SHL3 Final | tatcctactggagagagcaaggtcgctgttcaatacatgcACAGGATGTATA<br>TATCTGACACGTGCCTGGAGACTAGGGAGGAGAGACATTGGCGGTTACACCT<br>GGTGGGACAGCGCGTACGTGCGTTTAAGCGGTGCTAGAGCTGTCTACGACCA<br>ATTGAGCGGCCTCGGCACCGGGATTCTCCAGggcggccgcgtatagggtcca<br>tcacataagggatgaactc |
| 601-SHL6.5 Final | TatcctactggagagagcaaggtcgctgttcaatacatgcACAGGAGAGAGA<br>TAATCTGACACGTGCCTGGAGACTAGGGAGTAATCCCCTTGGCGGTTACACCT<br>GGTGGGACAGCGCGTACGTGCGTTTAAGCGGTGCTAGAGCTGTCTACGACCA<br>ATTGAGCGGCCTCGGCACCGGGATTCTCCAGggcggccgcgtatagggtcca<br>tcacataagggatgaactc |
| 601-SHL4.5 Final | TatcctactggagagagcaaggtcgctgttcaatacatgcACAGGATGTATA<br>TATCTGACACGTGCGAGAGAAATAGGGAGTAATCCCCTTGGCGGTTACACCT<br>GGTGGGACAGCGCGTACGTGCGTTTAAGCGGTGCTAGAGCTGTCTACGACCA<br>ATTGAGCGGCCTCGGCACCGGGATTCTCCAGggcggccgcgtatagggtcca<br>tcacataagggatgaactc |
| 601-SHL2.5 Final | tatcctactggagagagcaaggtcgctgttcaatacatgcACAGGATGTATA<br>TATCTGACACGTGCCTGGAGACTAGGGAGTAATCGAGAGAGAGGTTACACCT<br>GGTGGGACAGCGCGTACGTGCGTTTAAGCGGTGCTAGAGCTGTCTACGACCA<br>ATTGAGCGGCCTCGGCACCGGGATTCTCCAGggcggccgcgtatagggtcca<br>tcacataagggatgaactc |

| | |
|---|---|
| PCR template | pGEM3z-601 Addgene #26656 |
| 601 forward | TAT CC/iAmMC6T/ ACT GGA GAG AGC AAG GTC GC |
| SHL7 reverse | GTA CGC GCT GTC CCA CCA GGT GTA ACC GCC AAG GGG ATT ACT CCC TAG TCT CCA GGC ACG TGT CAG ATA T/iAmMC6T/T ACT CTC TCT GCA TGT ATT G |
| SHL5 reverse | GTA CGC GCT GTC CCA CCA GGT GTA ACC GCC AAG GGG ATT ACT CCC TAG TC/iAmMC6T/ CCA TCT CTC TGT CAG ATA T |
| SHL3 reverse | GTA CGC GCT GTC CCA CCA GGT GTA ACC GCC /iAmMC6T/AG GTC TCT CCT CCC TAG TC |
| SHL 6.5 reverse | GTA CGC GCT GTC CCA CCA GGT GTA ACC GCC AAG GGG ATT ACT CCC TAG TCT CCA GGC ACG TGT CAG A/iAmMC6T/A TCT CTC TCC TGT GCA T |
| SHL 4.5 reverse | GTA CGC GCT GTC CCA CCA GGT GTA ACC GCC AAG GGG ATT ACT CCC TA/iAmMC6T/ TTC TCT CGC ACG TGT CAG |
| SHL 2.5 reverse | GTA CGC GCT GTC CCA CCA GGT GTA ACC /iAmMC6T/CT CTC TCG ATT ACT CCC |
| Short fragment forward | AGT AAT CCC CTT GGC GGT TAC ACC TGG TGG GAC AGC GCG |
| Short fragment reverse | /5Biosg/GA GTT CAT CCC TTA TGT GAT GG |

*hsp70 DNA Constructs and Primers (2 GAF binding sites only)*

| | |
|---|---|
| hsp70 tandem final | TCATTTGTTTGGCAGAAAGAAAACTCTCAAAATTTCGTTGGCCGTTATAGCATATTCGTTTTGTGAGAGAGGGAGAGAGTACTATTGTTAACCCACAAGGTCGCACAGTAAACGGCACACTGTACGTGTTGCTTCGAGAGAGCGCGCcacctggtgtcgcgaaaagagcgccggagtataaatagag |
| hsp70 flipped final | TCATTTGTTTGGCAGAAAGAAAACTCTCAAAATTTCGTTGGCCGTTATAGCATATTCGTTTTGTGACTCTCCCTCTCTGTACTATTGTTAACCCACAAGGTCGCACAGTAAACGGCACACTGTACGTGTTGCTTCGAGAGAGCGCGCcacctggtgtcgcgaaaagagcgccggagtataaatagag |
| hsp70 Motif 2 Only final | TCATTTGTTTGGCAGAAAGAAAACTCTCAAAATTTCGTTGGCCGTTATAGCATATTCGTTTTGTCCAGCACTGTAATAGTACTATTGTTAACCCACAAGGTCGCACAGTAAACGGCACACTGTACGTGTTGCTTCGAGAGAGCGCGCcacctggtgtcgcgaaaagagcgccggagtataaatagag |
| PCR template | GAACGGGACAGGATACTTCCGATATCTCATTTGTTTGGCAGAAAGAAAACTCTCAAAATTTCGTTGGCCGTTATAGCATATTCGTTTTGTGACTCTCCCTCTCTGTACTATTGTTAACCCACAAGGTCGCACAGTAAACGGCACACTGTACGTGTTGCTTCGAGAGAGCGCGCCACCTGGTGTCGCGAAAAGAGCGCCGGAGTATAAATAGAGGATATCACCTCCCACTTACCTGTATT |
| Forward 1 (flipped) | TCA TTT GTT TGG CAG AAA GAA AAC TCT CAA AAT TTC GTT GGC CGT TAT AGC ATA TTC GTT TTG TGA CTC TCC CTC TCT G/iAmMC6T/A CTA TTG TTA AC |
| Forward 2 (tandem) | TCA TTT GTT TGG CAG AAA GAA AAC TCT CAA AAT TTC GTT GGC CGT TAT AGC ATA TTC GTT TTG TGA GAG AGG GAG AGA G/iAmMC6T/A CTA TTG TTA AC |
| Forward 3 (Motif 2 Only) | TCA TTT GTT TGG CAG AAA GAA AAC TCT CAA AAT TTC GTT GGC CGT TAT AGC ATA TTC GTT TTG TCC AGC ACT GTA ATA G/iAmMC6T/A CTA TTG TTA AC |
| Reverse | CTC CGG CGC TCT TTT CGC GAC ACC AGG TGG CGC GCT CTC TCG AAG C/iAmMC6T/A CAC GTA CAG |
| Short strand 1 | /5Phos/GT GTC GCG AAA AGA GCG CCG GAG TAT AAA TAG AG |
| Short strand 2 | /5Biosg/CT CTA TTT ATA CTC CGG CGC TCT TTT CGC GAC ACC AG |

Native *hsp70* NCP DNA

gtgTCATTTGTTTGGCAGAAAGAAAACTCGAGAAATTTCTCTGGCCGTTATTCTCTATTCGTTTTGTGACTCTCCCTCTCTGTACTATTGCTCTCTCACTCTGTCGCACAGTAAACGGCACACTGTTCTCGTTGCTTCGAGAGAGCGCGCctcgaa-Cy5
