## Supplementary Note for "GAGA zinc finger transcription factor searches chromatin by 1D-3D facilitated diffusion"

### Model Setup for GAF 1D Diffusion

Our single-molecule FRET data show GAF-DBD performs 1D diffusion on the linear DNA to search for the cognate motif. GAF-DBD is labeled with the FRET donor Cy3 (shown in green), and two motifs on the DNA are labeled with the FRET acceptors Cy7 (blue) and Cy5 (red), respectively. When GAF-DBD binds on the DNA, under Cy3 excitation, the 3 fluorophores will show different intensity signatures depending on GAF-DBD's position on the DNA. As the GAF-DBD moves along the DNA, the 3 fluorophores' intensities over time are recorded as the fluorescence intensity time trajectory. These 3-color FRET values can occupy an approximately continuous space, but we will map them to 6 hidden states, one where GAF-DBD is in solution, and 5 on the linear DNA. The states on DNA are labeled with ① to ⑤ (Fig S1A). Position ① is located at the far left, where we expect high Cy3 intensity, and low Cy5 and Cy7 intensities. Position ② is located at the motif labeled with Cy7, so we expect high Cy7 intensity, medium Cy3 intensity, and low Cy5 intensity. Position ③ is located between ② and ④, where all 3 fluorophores show medium intensities. Position ④ is located on the motif labeled with Cy5, exhibiting high Cy5 intensity, medium Cy3 intensity, and low Cy7 intensity. Position ⑤ is located at the far right, showing high Cy3 intensity and low Cy5 and Cy7 intensity. Besides the 5 states on the DNA, we also define state ∅ as freely diffusing GAF-DBD in solution, where all 3 fluorophores show low intensities. A long trajectory often contains all 6 states, as shown in Fig S1B and C.

According to the above description of the 6 states, the green fluorophore has only three different fluorescence intensity values, so do the blue and red fluorophores,

$$\vec{u}^g = \left[u_h^g, u_m^g, u_l^g\right]^T$$
$$\vec{u}^b = [u_h^b, u_m^b, u_l^b]^T \ , \qquad (1)$$
$$\vec{u}^r = [u_h^r, u_m^r, u_l^r]^T$$

where the superscript $g$ refers to the green fluorophore, $b$ refers to the blue fluorophore, and $r$ refers to the red fluorophore; the subscript $h$ refers to the high intensity, $m$ refers to the medium intensity, and $l$ refers to the low intensity. Then, the total 6 states of the system can be written as

$$U = \begin{bmatrix} u_l^g & u_h^g & u_m^g & u_m^g & u_m^g & u_h^g \\ u_l^b & u_l^b & u_h^b & u_m^b & u_l^b & u_l^b \\ u_l^r & u_l^r & u_l^r & u_m^r & u_h^r & u_l^r \end{bmatrix} \begin{matrix} \longrightarrow \text{Cy3 (green-color)} \\ \longrightarrow \text{Cy7 (blue-color)} \\ \longrightarrow \text{Cy5 (red-color)} \end{matrix} \qquad (2)$$
$$\begin{matrix} \downarrow & \downarrow & \downarrow & \downarrow & \downarrow & \downarrow \\ \varnothing & ① & ② & ③ & ④ & ⑤ \end{matrix}$$

Each column of the above matrix, $\vec{U}_i$ $(i = 0 \dots 5)$ represents each state of the system and corresponds to one specific position on DNA. The 3 rows represent the 3 fluorophores' fluorescence intensities respectively. Eq (2) clearly shows the 6 states are not fully independent, instead they are partially coupled because some elements should have the same values. The couplings in Eq (2) not only determines relations between each system state, but also guarantees that the 6 positions are spatially ordered as in FigS1A. With the couplings, the system $U$ has in total 9 values for optimization, which are the 3 fluorophores' intensities, $\vec{u}^g$, $\vec{u}^b$, and $\vec{u}^r$.

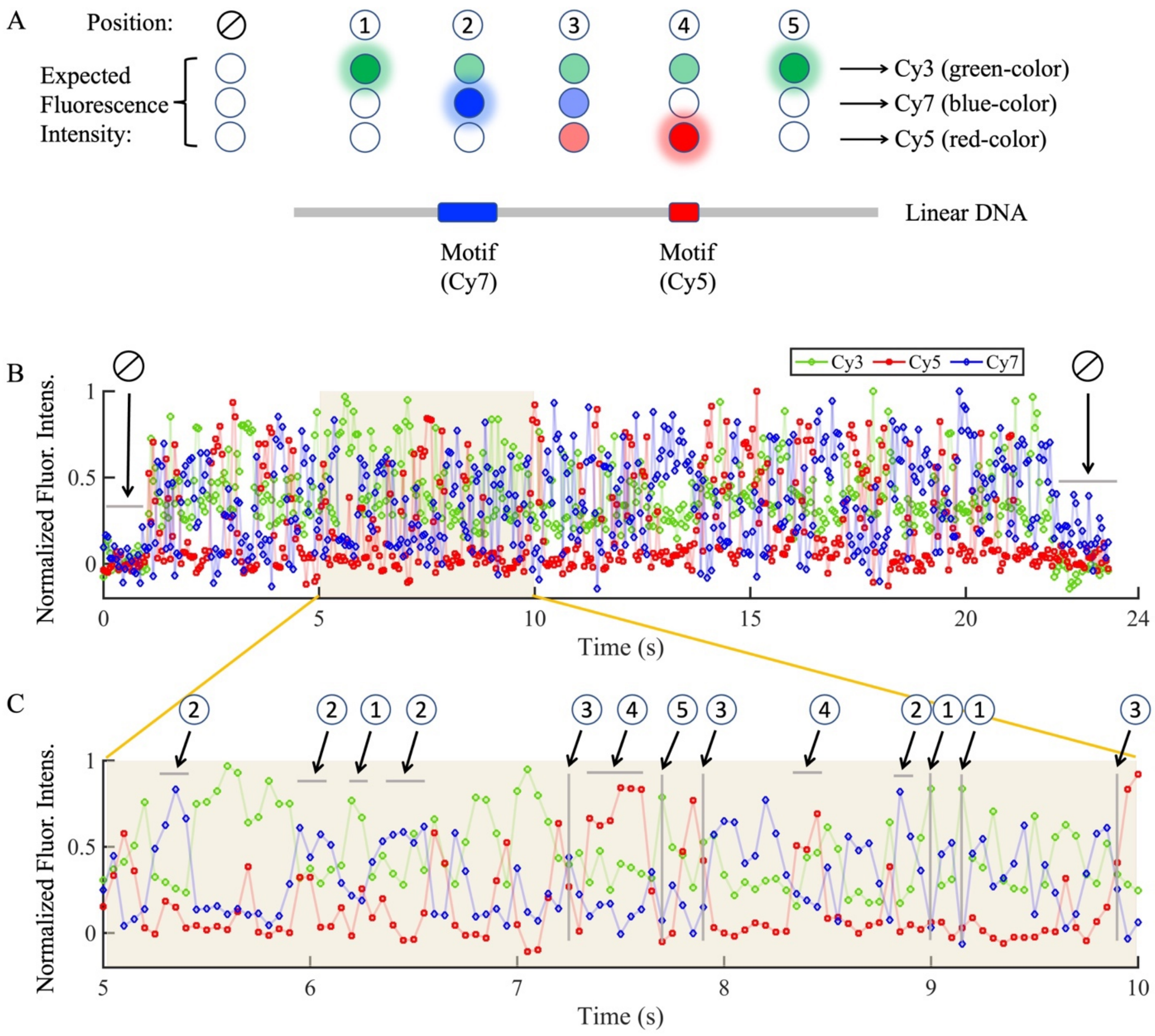


Figure S1. GAF-DBD positions on linear DNA and the relevant fluorescence intensity time trajectory. (A) Six selected positions of GAF-DBD on linear DNA. (B) One time trajectory. Markers are experimental data connected by light lines in time order. The green data is the Cy3 intensity, the blue data is the Cy7 intensity, and the red data is the Cy5 intensity. The beginning and the ending of the trajectory both represent the state where GAF-DBD is in solution and the corresponding position is $\varnothing$. (C) A segment of the time trajectory in (B). Several representative positions are highlighted that illustrate the expected FRET emissions given their location on the DNA.

### Hidden Markov Model for the Analysis of 3-Color FRET Data

From our FRET data, we interpret the fluorescence signal over time as a series of well-defined states. An appropriate model to quantify the probabilities of these states and the transitions between them is the Hidden Markov Model (HMM), which has previously been applied to analyze 2-color FRET data [1-3]. We will introduce the HMM framework and the constraints for the GAF-DBD model, followed by our modifications to the optimization algorithm.

The time series of FRET data consists of state-to-state transitions of the system. Because of the data noise, the underling state at each time and the specific state transition events are obscured which makes the process a hidden Markov process.

Consider our system here with 3 types of fluorophores and has 6 possible combinations of their intensity levels. At each time $t$, the observed FRET data is composed of 3 elements, denoted by a vector $\vec{x}_t$. The hidden state $s_t$ at time $t$ can be any one of the 6 states.

The probability that the observed data $\vec{x}_t$ comes from the state $s_t$ is the emission probability $p(\vec{x}_t|s_t)$, which is modeled by a Gaussian mixture model. The emission probability is a function of the mean value $\vec{U}_i$ and covariance $\sigma_i$ of the Gaussian distribution for each state, $i \in \{0, 1, 2, 3, 4, 5\}$. The emission probability is also a function of the state number $S$. $S$ is generally an unknown variable, but here we set $S$ equals to 6. The probability of the entire FRET trajectory is obtained by multiplying the product of the emission probability and the transition probability matrix $A(i, j)$, where $i, j \in \{0, 1, 2, 3, 4, 5\}$,

$$P = \prod_{t=1}^{N} [p(\vec{x}_t|s_t) \cdot A(s_{t-1}, s_t)], \quad (3)$$

where $N$ is the total number of time points. $P$ is also called the likelihood function. Then, one needs to find the specific values of $\vec{U}_i^*$, $\sigma_i^*$ and $A^*$ to maximize the likelihood function,

$$P^* = \max_{\vec{U}_i^*, \sigma_i^*, A^*} (P). \quad (4)$$

The multivariant optimization problem described by Eq (4) can be solved by many numerical algorithms, among which the expectation-maximization (EM) algorithm is the most efficient one [4]. Once the solution of Eq (4) is obtained, one can use the Viterbi algorithm to find the most likely sequence of states at each time point [4].

**Constraint EM Algorithm**

The traditional EM algorithm does not include the couplings among system states because the M step uses $\partial P/\partial \vec{U}_i = 0$ to update each state $\vec{U}_i$, under the assumption that all system states are independent from each other. Our analysis shows traditional EM generates 18 different values for system states $U$, and therefore fails to properly constrain locations of states along the DNA. When we used HMM with the traditional EM algorithm to analyze 30 FRET trajectories with at least 70 time points, only 30% of the trajectories passably gave the correct corresponding positions to the 6 states. Increasing the number of the states and positions did not significantly improve the results due to over-fitting.

Therefore, we developed a new method to analyze the FRET data. Our method is based on the traditional EM algorithm with the extra constraints that describe the couplings among the system states. Each state of the system is a combination of the 3 fluorescence intensity levels, which can be written as:

$$\vec{U}_i = M_i^g \vec{u}^g + M_i^b \vec{u}^b + M_i^r \vec{u}^r, \quad (5)$$

where $M$ is a coefficient matrix with the dimension 3×3, and $\vec{u}^g$, $\vec{u}^b$ and $\vec{u}^r$ are the target values of the 3 fluorophores as in Eq (1). $M$ determines which element of $\vec{u}^g$, $\vec{u}^b$ or $\vec{u}^r$ is selected to compose the system state. Therefore, $M$ is also called the selection-matrix. According to the expected 6 states in Eq (2), we get all the 18 selection-matrices,

$$M_0^g = \begin{bmatrix} 0 & 0 & 1 \\ 0 & 0 & 0 \\ 0 & 0 & 0 \end{bmatrix}, M_0^b = \begin{bmatrix} 0 & 0 & 0 \\ 0 & 0 & 1 \\ 0 & 0 & 0 \end{bmatrix}, M_0^r = \begin{bmatrix} 0 & 0 & 0 \\ 0 & 0 & 0 \\ 0 & 0 & 1 \end{bmatrix}$$

$$M_1^g = \begin{bmatrix} 1 & 0 & 0 \\ 0 & 0 & 0 \\ 0 & 0 & 0 \end{bmatrix}, M_1^b = \begin{bmatrix} 0 & 0 & 0 \\ 0 & 0 & 1 \\ 0 & 0 & 0 \end{bmatrix}, M_1^r = \begin{bmatrix} 0 & 0 & 0 \\ 0 & 0 & 0 \\ 0 & 0 & 1 \end{bmatrix}$$

$$M_2^g = \begin{bmatrix} 0 & 1 & 0 \\ 0 & 0 & 0 \\ 0 & 0 & 0 \end{bmatrix}, M_2^b = \begin{bmatrix} 0 & 0 & 0 \\ 1 & 0 & 0 \\ 0 & 0 & 0 \end{bmatrix}, M_2^r = \begin{bmatrix} 0 & 0 & 0 \\ 0 & 0 & 0 \\ 0 & 0 & 1 \end{bmatrix} \tag{6}$$

$$M_3^g = \begin{bmatrix} 0 & 1 & 0 \\ 0 & 0 & 0 \\ 0 & 0 & 0 \end{bmatrix}, M_3^b = \begin{bmatrix} 0 & 0 & 0 \\ 0 & 1 & 0 \\ 0 & 0 & 0 \end{bmatrix}, M_3^r = \begin{bmatrix} 0 & 0 & 0 \\ 0 & 0 & 0 \\ 0 & 1 & 0 \end{bmatrix}$$

$$M_4^g = \begin{bmatrix} 0 & 1 & 0 \\ 0 & 0 & 0 \\ 0 & 0 & 0 \end{bmatrix}, M_4^b = \begin{bmatrix} 0 & 0 & 0 \\ 0 & 0 & 1 \\ 0 & 0 & 0 \end{bmatrix}, M_4^r = \begin{bmatrix} 0 & 0 & 0 \\ 0 & 0 & 0 \\ 1 & 0 & 0 \end{bmatrix}$$

$$M_5^g = \begin{bmatrix} 1 & 0 & 0 \\ 0 & 0 & 0 \\ 0 & 0 & 0 \end{bmatrix}, M_5^b = \begin{bmatrix} 0 & 0 & 0 \\ 0 & 0 & 1 \\ 0 & 0 & 0 \end{bmatrix}, M_5^r = \begin{bmatrix} 0 & 0 & 0 \\ 0 & 0 & 0 \\ 0 & 0 & 1 \end{bmatrix}$$

In Eq (5) and Eq (6), the system $U$ is not the independent parameter for the optimization, instead, $\vec{u}^g$, $\vec{u}^b$ and $\vec{u}^r$ are the independent parameters which have 9 values in total.

In the maximization step of the constraint EM algorithm, to make the system likelihood maximized, values of $\vec{u}^g$, $\vec{u}^b$ and $\vec{u}^r$ should satisfy $\partial P/\partial\vec{u}^g = 0$, $\partial P/\partial\vec{u}^b = 0$ and $\partial P/\partial\vec{u}^r = 0$, respectively, which can be rewritten as

$$\frac{\partial P}{\partial \vec{u}^g} = \sum_{i=0}^{5} \left[\frac{\partial U_i}{\partial (\vec{u}^g)^T}\right]^T \cdot \frac{\partial P}{\partial \vec{U}_i} = \sum_{i=0}^{5} \left(M_i^g\right)^T \cdot \frac{\partial P}{\partial \vec{U}_i} = 0$$

$$\frac{\partial P}{\partial \vec{u}^b} = \sum_{i=0}^{5} \left[\frac{\partial U_i}{\partial \left(\vec{u}^b\right)^T}\right]^T \cdot \frac{\partial P}{\partial \vec{U}_i} = \sum_{i=0}^{5} \left(M_i^b\right)^T \cdot \frac{\partial P}{\partial \vec{U}_i} = 0 \tag{7}$$

$$\frac{\partial P}{\partial \vec{u}^r} = \sum_{i=0}^{5} \left[\frac{\partial U_i}{\partial (\vec{u}^r)^T}\right]^T \cdot \frac{\partial P}{\partial \vec{U}_i} = \sum_{i=0}^{5} (M_i^r)^T \cdot \frac{\partial P}{\partial \vec{U}_i} = 0$$

The term of $\partial P/\partial\vec{U}_i$ can be found in the traditional EM algorithm, like the Eq 9.16 in [4],

$$\frac{\partial P}{\partial \vec{U}_i} = -\sum_{n=1}^{N} \gamma_{ni} \sigma_i \left(\vec{x}_n - \vec{U}_i\right) \tag{8}$$

where $\gamma$ is the posterior probability, and σ is the covariance of the Gaussian distribution which is a symmetric matrix [4]. By substituting Eq (8) and Eq (5) into Eq (7), we can get

$$A_{11} \cdot \vec{u}^g + A_{12} \cdot \vec{u}^b + A_{13} \cdot \vec{u}^r = \vec{C}_1$$

$$A_{21} \cdot \vec{u}^g + A_{22} \cdot \vec{u}^b + A_{23} \cdot \vec{u}^r = \vec{C}_2 \tag{9}$$

$$A_{31} \cdot \vec{u}^g + A_{32} \cdot \vec{u}^b + A_{33} \cdot \vec{u}^r = \vec{C}_3$$

or rewritten as

$$\begin{bmatrix} A_{11} & A_{12} & A_{13} \\ A_{21} & A_{22} & A_{23} \\ A_{31} & A_{32} & A_{33} \end{bmatrix} \begin{bmatrix} \vec{u}^g \\ \vec{u}^b \\ \vec{u}^r \end{bmatrix} = \begin{bmatrix} \vec{C}_1 \\ \vec{C}_2 \\ \vec{C}_3 \end{bmatrix},$$

where

$A_{11} = \sum_{i=0}^{5} \left(M_i^g\right)^T \sigma_i\, M_i^g \left(\sum_{n=1}^{N} \gamma_{ni}\right)$,

$A_{12} = \sum_{i=0}^{5} \left(M_i^g\right)^T \sigma_i\, M_i^b \left(\sum_{n=1}^{N} \gamma_{ni}\right)$,

$A_{13} = \sum_{i=0}^{5} \left(M_i^g\right)^T \sigma_i\, M_i^r \left(\sum_{n=1}^{N} \gamma_{ni}\right)$,

$A_{21} = \sum_{i=0}^{5} \left(M_i^b\right)^T \sigma_i\, M_i^g \left(\sum_{n=1}^{N} \gamma_{ni}\right)$,

$A_{22} = \sum_{i=0}^{5} \left(M_i^b\right)^T \sigma_i\, M_i^b \left(\sum_{n=1}^{N} \gamma_{ni}\right)$,

$A_{23} = \sum_{i=0}^{5} \left(M_i^b\right)^T \sigma_i\, M_i^r \left(\sum_{n=1}^{N} \gamma_{ni}\right)$,

$A_{31} = \sum_{i=0}^{5} (M_i^r)^T \sigma_i\, M_i^g \left(\sum_{n=1}^{N} \gamma_{ni}\right)$,

$A_{32} = \sum_{i=0}^{5}(M_i^r)^T \sigma_i \, M_i^b (\sum_{n=1}^{N} \gamma_{ni})$ ,
$A_{33} = \sum_{i=0}^{5}(M_i^r)^T \sigma_i \, M_i^r (\sum_{n=1}^{N} \gamma_{ni})$ ,
$\vec{C}_1 \quad = \sum_{i=0}^{5}\left(M_i^g\right)^T \sigma_i \, (\sum_{n=1}^{N} \gamma_{ni}\vec{x}_n)$ ,
$\vec{C}_2 \quad = \sum_{i=0}^{5}\left(M_i^b\right)^T \sigma_i \, (\sum_{n=1}^{N} \gamma_{ni}\vec{x}_n)$ ,
$\vec{C}_3 \quad = \sum_{i=0}^{5}(M_i^r)^T \sigma_i \, (\sum_{n=1}^{N} \gamma_{ni}\vec{x}_n)$ .
$A$ matrices in Eq (9) have the dimension 3×3, and $\vec{C}$ vectors have the dimension 3×1. The Eq (9) is a system of linear equations with a 9×9 coefficient matrix, which can be solved easily by a mathematical software such as MATLAB, to obtain values of $\vec{u}^g$, $\vec{u}^b$ and $\vec{u}^r$.

In the constraint EM algorithm, we apply the following three steps,

1. E step: It is the same as the traditional E step.
   Calculate the posterior probability and forward probability.
2. M step: According to Eq (9), calculate all $A$s and $\vec{C}$s, then solve for $\vec{u}^g$, $\vec{u}^b$ and $\vec{u}^r$.
   According to Eq (5), update each of the system state $\vec{U}_i$.
   Update the covariance and the transition matrix as the traditional M step.
3. Repeat 1-2 steps until parameters converge.

We programmed the HMM with the constraint EM algorithm in MATLAB, and the code is open source to the public: https://github.com/Yibenfu/HMM_3colorFRET.

**HMM with the Constraint EM Algorithm Successfully Analyzes 3-Color FRET Data**

The constraint EM algorithm effectively classifies our FRET data (Fig S2). The optimization gives 6 states, and each fluorophore has 3 distinct intensity values (Fig S3C). These 6 states can be unambiguously recognized to correspond to the 6 expected positions as illustrated in Fig S1A. According to the predicted hidden states in Fig S2A (higher panel), we can obtain the state sequence of the system which shows how GAF-DBD diffuses on the linear DNA, Fig S2A (lower panel).

We applied the HMM with the constraint EM algorithm to analyze 30 FRET trajectories with at least 70 time points, and this time 80% are successfully assigned. An assignment is successfully if the states and transitions are consistent with the expected fluorescence intensities when GAF-DBD dwells at the assigned position on the DNA. The other 7 trajectories fail to converge due to anomalies in the FRET data that we attribute to rare events such as transient appearance of multiple Cy3-labeled GAF-DBD molecules (Fig S3A). For this type of trajectories with few and isolated abnormal FRET data, we can adjust initial parameter estimates prior to the optimization (specifically, increasing estimated state covariances), which typically allows the HMM to converge. Alternatively, sustained irregular signals can be removed prior to optimization (Fig S3B). In principle, by adjusting initial inputs or deleting the abnormal data, we can apply HMM with the constraint EM algorithm to analyze all our FRET trajectories.

In general, this method outperforms the Relative Intensity Algorithm (see Methods) in capturing transient nonspecific states (Fig S4) from 3-color single-molecule trajectories. This is mainly because RI assigns a nonspecific state only when both Cy5 and Cy7 fluorescence is close to background, and thus would miss transient nonspecific states (< 35-50 ms exposure time) where the total signal per frame is weak but significant in Cy5 and/or Cy7 fluorescence and relatively high in Cy3 fluorescence. On the other hand, by considering the fluorescence signature of all 3 fluorophores, HMM is more sensitive to transient nonspecific states.

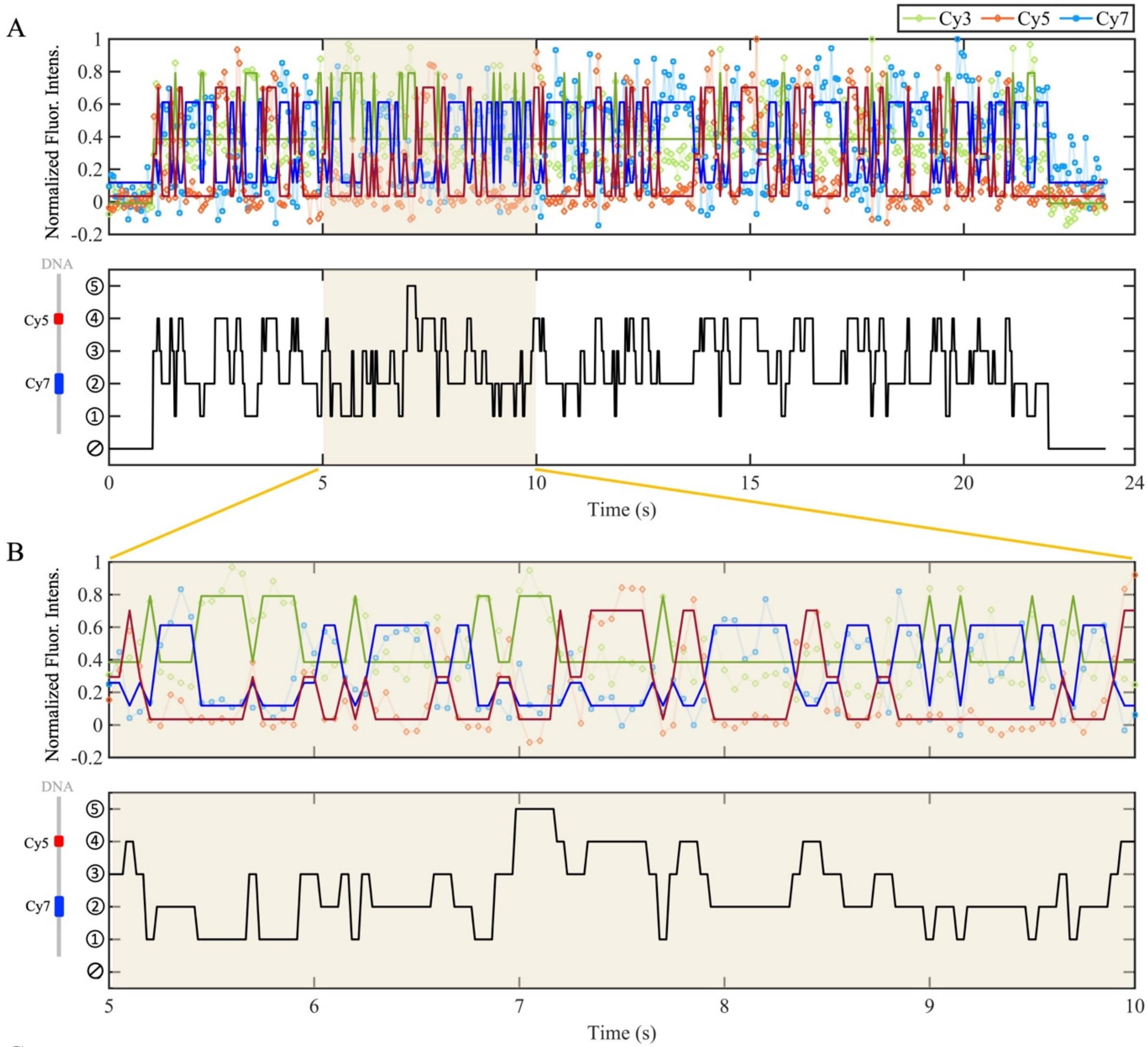


C

$$\begin{aligned} \text{Cy3 FRET: } \vec{u}^{g} &= [0.79 \quad 0.39 \quad -0.01]^{T} \\ \text{Cy7 FRET: } \vec{u}^{b} &= [0.61 \quad 0.26 \quad 0.12]^{T} \\ \text{Cy5 FRET: } \vec{u}^{r} &= [0.70 \quad 0.29 \quad 0.03]^{T} \end{aligned} \xrightarrow[\text{by Eq(5)}]{\text{build the state}} U = \begin{bmatrix} -0.01 & 0.79 & 0.39 & 0.39 & 0.39 & 0.79 \\ 0.12 & 0.12 & 0.61 & 0.26 & 0.12 & 0.12 \\ 0.03 & 0.03 & 0.03 & 0.29 & 0.70 & 0.03 \end{bmatrix}$$

↓ ∅ ↓ ① ↓ ② ↓ ③ ↓ ④ ↓ ⑤

Figure S2. FRET analysis by HMM with the constraint EM algorithm. (A) (Higher panel) Markers are experimental data connected by light lines in time order, which are the same as FigS1B. The solid thick lines are the predicted hidden state values for the FRET emission. (Lower panel) From the hidden state values shown in the upper panel we can assign each time-point to the most-likely state of GAF-DBD position on DNA using Viterbi, as shown here. This state sequence thus reports on the diffusion of the GAF-DBD between states. (B) A zoom-in of the same data from (A). (C) Optimized parameters for the mean FRET emissions for all six states of the system, as defined from the optimized mean fluorescence intensities for each fluorophore.

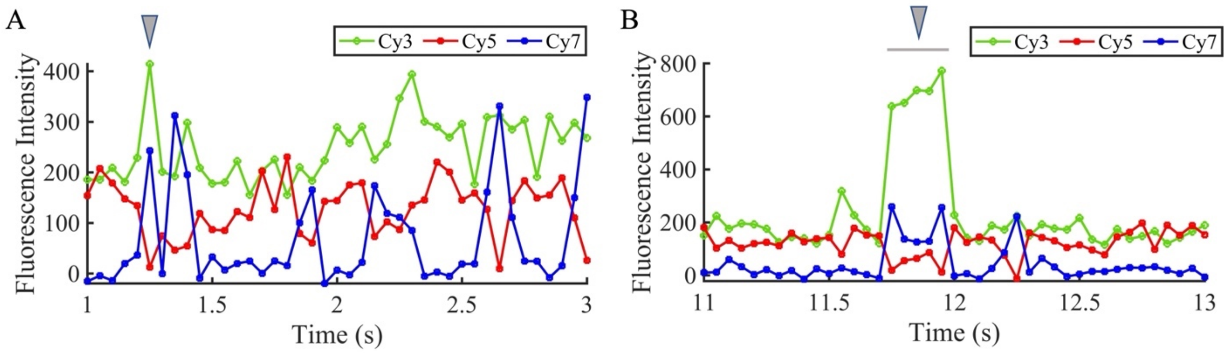


Figure S3. Examples of fluorescence intensity time trajectories with data indicating rare events that are not well-described by the six-state model. (A) Example of transient encounters between two GAF-DBD on DNA. At the moment ~1.25s, it shows the green fluorophore has a high intensity, meanwhile the blue fluorophore also has a high (at least medium) intensity. This can't happen according to our above six-state model, because the green fluorophore should not hold a high intensity when it inspires the blue fluorophore. We suspect that the high green signal originates from more than one GAF-DBD simultaneously and transiently appearing near the motif with the blue fluorophore, resulting in both green and blue fluorescence. (B) Example of persistently co-localized GAF-DBDs on DNA. At the time 11.8–12s, the green fluorophore shows extremely high fluorescence intensity, which can be similarly owing to multiple GAF-DBDs appearing on DNA at the same time. However, in this trajectory, the abnormal data is not isolated, instead it is consistent for a time interval ~0.2s. For this type of trajectories, the best operation before running the HMM analysis is to delete these abnormal data.

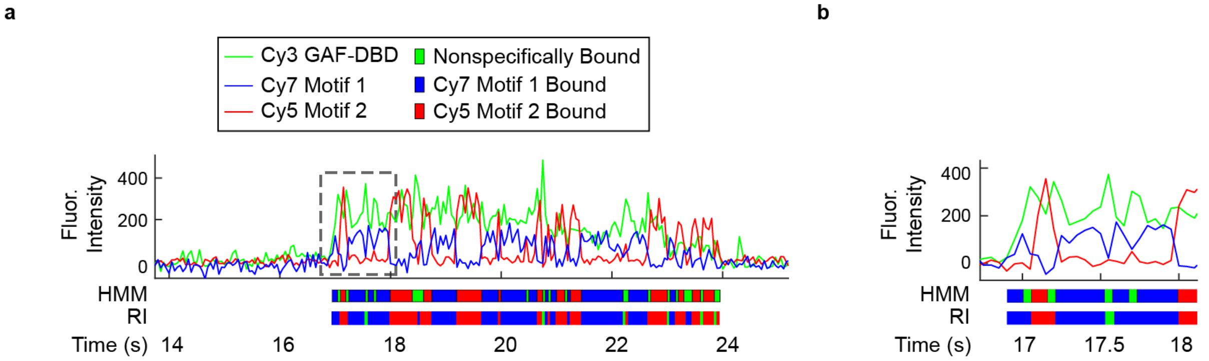


Figure S4. HMM with Constraint EM Algorithm (HMM) better captures transient nonspecific states compared to the Relative Intensity Algorithm.

### Reference


1. Eddy, S.R., *What is a hidden Markov model?* Nature Biotechnology, 2004. **22**(10): p. 1315-1316.
2. McKinney, S.A., C. Joo, and T. Ha, *Analysis of single-molecule FRET trajectories using hidden Markov modeling.* Biophysical journal, 2006. **91**(5): p. 1941-1951.
3. Bronson, J.E., et al., *Learning rates and states from biophysical time series: a Bayesian approach to model selection and single-molecule FRET data.* Biophysical journal, 2009. **97**(12): p. 3196-3205.
4. Bishop, C.M., *Pattern recognition and machine learning*. Information science and statistics. 2006, New York: Springer. xx, 738 p.
